# A Mechanism for Epithelial-Mesenchymal Heterogeneity in a Population of Cancer Cells

**DOI:** 10.1101/592691

**Authors:** Shubham Tripathi, Herbert Levine, Mohit Kumar Jolly

## Abstract

Epithelial-mesenchymal heterogeneity, wherein cells within the same tumor can exhibit an epithelial, a mesenchymal, or one or more hybrid epithelial-mesenchymal phenotype(s), has been observed across cancer types and implicated in metastatic aggressiveness. Here, we have used computational modeling to show that this heterogeneity can emerge from the noise in the partitioning of RNAs and proteins among the daughter cells during cancer cell division. Our model captures the population-level behavior of murine prostate cancer cells, the hysteresis in the dynamics of epithelial-mesenchymal plasticity, and how hybrid phenotype-promoting factors alter the phenotypic composition of a population. We further used the model to describe the implications of heterogeneity for therapeutics. By linking the dynamics of an intracellular regulatory circuit to the phenotypic composition of a population, the study contributes towards understanding how non-genetic heterogeneity can be generated and propagated from a small, homogeneous population, and towards therapeutic targeting of cancer cell heterogeneity.

## Introduction

A tumor can contain cancer cells exhibiting multiple, distinct phenotypes. Subpopulations of cells exhibiting a stem cell-like phenotype have been reported in multiple cancer types including leukemia (Lapidot et al., 1994), breast cancer (Al-Hajj et al., 2003), colorectal cancer (O’Brien et al., 2007; Ricci-Vitiani et al., 2007), brain cancer (Singh et al., 2004), and prostate cancer (Collins et al., 2005). Tumor cell populations in triple-negative breast cancer can consist of luminal, basal, immunomodulatory, mesenchymal, and stem-like cells (Lehmann et al., 2011), while those in small cell lung cancer are composed of neuroendocrine and non-neuroendocrine cells (Calbo et al., 2011; Udyavar et al., 2016). Intra-tumoral heterogeneity has been associated with poor prognosis across cancer types (Stanta and Bonin, 2018). Cancer cells within a tumor can exhibit different phenotypes in spite of carrying the same genetic alterations, indicating that a non-genetic mechanism underlies the phenotypic heterogeneity. Further, these cells can switch between different phenotypic states, either spontaneously or in response to specific cues. Spontaneous switching between luminal, basal, and stem-like states has been reported in breast cancer cell lines (Gupta et al., 2011) while androgen-deprivation therapy has been shown to promote transition to a neuroendocrine phenotype in prostate cancer (Hirano et al., 2004; Wright et al., 2003). Cancer cells with different phenotypes can exhibit different sensitivities to various drugs and therapeutic regimens (Risom et al., 2018; Wooten et al., 2018). Therefore, non-genetic mechanisms of phenotypic heterogeneity and plasticity in cancer cells are a fundamental challenge to anti-cancer therapies and an understanding of such mechanisms is essential to the development of effective anti-cancer treatments.

A canonical example of non-genetic heterogeneity is epithelial-mesenchymal plasticity (EMP). It involves two reversible processes, epithelial-mesenchymal transition (EMT) and mesenchymal-epithelial transition (MET). Via EMT, cancer cells, to varying extents, can lose epithelial characteristics such as cell-cell adhesion and apico-basal polarity, and acquire mesenchymal features which allow cancer cells to migrate effectively and invade. The reverse change is observed during MET-cells lose their migratory freedom and re-acquire epithelial hallmarks including expression of junctional proteins (Nieto et al., 2016). Recent studies have shown that cancer cells can also stably exist in a hybrid epithelial-mesenchymal state wherein they co-express epithelial and mesenchymal markers (Jolly et al., 2016; Pastushenko et al., 2018). Cancer cells within a solid tumor exhibit widespread heterogeneity in the expression of epithelial and mesenchymal markers and can exhibit an epithelial (E), a mesenchymal (M), or one or more hybrid epithelial-mesenchymal (hybrid E / M) phenotype(s) (Hong et al., 2018; Kim et al., 2013; Pereira et al., 2015; Stylianou et al., 2018).

By allowing the tumor cells to acquire migratory traits for dissemination to distant organs, EMP plays a key role in the metastatic spread of solid tumors. The disseminated cells can then re-acquire epithelial traits including cell-cell adhesion to establish a tumoral mass at the new organ site. EMP has further been implicated in the triggering of stemness programs in cancer cells (Bocci et al., 2019; Mani et al., 2008), evasion of the host immune response (Kudo-Saito et al., 2009), and emergence of resistance to anti-cancer therapies (Fischer et al., 2015; Marín-Aguilera et al., 2014; Zheng et al., 2015). Hybrid E / M cells play a key role in the collective dissemination of tumor cells as clusters, an aggressive mechanism of cancer metastasis (Jolly et al., 2015). Further, EMP-associated phenotypes differ in their tumor-seeding abilities (Grosse-Wilde et al., 2018; Neelakantan et al., 2017) and in their sensitivity to drugs (Creighton et al., 2009; Tièche et al., 2018). Understanding the mechanisms driving EMP will thus be a critical step in the development of more effective anti-cancer therapies.

The signaling pathways and the various transcription factors, micro-RNAs, and environmental stimuli driving EMP are well characterized (Nieto et al., 2016). The population-level dynamics of EMP, on the other hand, has only recently come into focus. Ruscetti *et al.* showed, in a mouse model of prostate cancer, that cells exhibiting a single EMP-associated phenotype can spontaneously generate a population with all three phenotypes over a period of time (Ruscetti et al., 2016). Celià-Terrassa *et al.* observed hysteresis in the dynamics of EMP in multiple normal and cancerous mammary epithelial cell lines and reported a role for hysteretic dynamics in the metastatic ability of cancer cells (Celià-Terrassa et al., 2018). Risom *et al.* showed that drugs targeting specific cellular pathways alter the transitions between epithelial-like and mesenchymal-like states in breast cancer cell lines, thereby establishing phenotypic plasticity as a therapeutically targetable property (Risom et al., 2018). The mechanism responsible for the phenotypic heterogeneity and plasticity in cancer cells, however, remains uncharacterized. Some studies have described phenomenological models of phenotypic plasticity in cancer cells (Chapman et al., 2019; Gupta et al., 2011; Risom et al., 2018). While predictions from these models fit the experimental data reported in the respective studies, such models lack detailed biomolecular mechanistic bases. This limitation makes the use of such phenomenological models difficult for obtaining predictions directly useful in designing anti-cancer therapies.

Multiple mechanisms can drive phenotypic heterogeneity in a population. Apart from stochastic changes in the phenotypes of cells in a population due to intrinsic noise, heterogeneity can also arise from variable strengths of regulatory interactions in the cells in a population. The regulatory circuits in different cells can then exhibit distinct dynamics in response to the same external cues (Huang et al., 2017). Another mechanism by which heterogeneity can emerge is cell-cell communication. A prominent example is Notch-Delta-Jagged signaling wherein cells in a population can exhibit a sender phenotype, a receiver phenotype, or a hybrid sender-receiver phenotype (Boareto et al., 2016). Here, we focus on the emergence of heterogeneity via the first mechanism i.e. stochastic changes in the phenotypes of cells in response to noise.

Stochastic changes in cell phenotype, in general, require a mechanism to generate noise and a mechanism to stabilize the decision reached in response to it (Losick and Desplan, 2008). In both prokaryotes and eukaryotes, noise can arise from the inherent stochasticity of the transcription process. A typical case is the emergence of antibiotic-resistant persister cells in bacterial populations (Balaban et al., 2004). Another source of noise is the random partitioning of parent cell biomolecules among the daughter cells at the time of cell division (Huh and Paulsson, 2011a). A key role for such noise in the generation of non-genetic heterogeneity has been proposed (Huh and Paulsson, 2011b). Fluctuations in the phenotype in response to noise are usually small and transient. Therefore, stochastic cell-phenotype changes also require a mechanism to amplify the fluctuations in the phenotype in response to noise. This can be achieved if the underlying response mechanism exhibits multi-stability. In such a scenario, even small fluctuations added to a steady state can cause the cell to transition to a new stable steady state. Cancer cells divide uncontrollably, making random partitioning of biomolecules during cell division a significant source of noise. The regulatory circuit governing EMP has been shown to exhibit tri-stability (Lu et al., 2013). Thus, both the requirements for stochastic cell-phenotype changes are satisfied for a population of cancer cells exhibiting EMP.

We show that in a population of cells each carrying a copy of the EMP regulatory circuit, phenotypic heterogeneity can arise from the noise generated due to random partitioning of micro-RNAs, mRNAs, and transcription factors at the time of cell division. The temporal dynamics of fractions of different phenotypes in a population predicted by our model agrees with that observed for cells isolated from a mouse model of prostate cancer (Ruscetti et al., 2016), both in terms of the timescale over the phenotypic composition of the population changes and the phenotypic distribution under different initial conditions. Our model captures the relative stability of epithelial and mesenchymal populations and the ability of a hybrid E / M population to quickly generate a mixture of epithelial and mesenchymal cells. We also used the model to describe hysteresis in the dynamics of EMP and characterized how EMP modulators such as GRHL2 and ΔNP63α can alter the phenotypic composition of a population. Our model predicted that a suitable combination of an EMT promoter, such as retinoic acid, and an EMT inhibitor, such as TGF-*β*, can stabilize a population of hybrid E / M cells. Finally, we characterized how drugs targeting specific EMP-associated phenotypes alter the population behavior. We conclude that targeting a single phenotype is likely to have little attenuating influence on tumor size. Targeting epithelial and mesenchymal cells simultaneously is predicted to be the best regimen for anti-cancer therapy.

## Results

### Developing a Population-level Model of EMP Dynamics

Epithelial-mesenchymal plasticity (EMP) is modulated via the functional cooperation of multiple cell signaling pathways that can respond to both internal and external stimuli (Lamouille et al., 2014). These include TGF-*β* SMAD3, WNT-*β*-catenin, and Notch pathways. The activity of these pathways converges onto a core regulatory circuit shown in fig. 1 (A). Downstream targets of this circuit include cadherins, claudins, occludins, and metalloproteinases (Lamouille et al., 2014). Due to the central role of this circuit in modulating EMP (Lamouille et al., 2014; Nieto et al., 2016), we used this regulatory circuit to construct a model of epithelial-mesenchymal heterogeneity in cancer cells. The EMP regulatory circuit forms a ternary switch, the dynamics of which has been described previously (Lu et al., 2013). Briefly, the circuit can exhibit three distinct steady states in response to different levels of the *SNAI1* activator *I*_*sig*_. These stable steady states can be mapped to the three EMP-associated phenotypes: epithelial (E), mesenchymal (M), and hybrid epithelial-mesenchymal (hybrid E / M) (fig. 1 (B)). We considered a population of cancer cells where each cell is carrying a copy of this regulatory circuit. The kinetic parameters governing the behavior of the EMP regulatory circuit have been described previously (Lu et al., 2013). The parameters are such that the circuit operates in the tri-stable region. This choice of parameters allowed us to explore the behavior of all three EMP-associated phenotypes and is suited for describing experiments characterizing epithelial-mesenchymal heterogeneity. The dynamics of the EMP regulatory circuit within each cell can be simulated independent of the other cells in the population. Each cell can then be assigned one of the three phenotypes based on the expression level of ZEB1 mRNA inside the cell. In the absence of competition (or, equivalently, in the presence of an infinite supply of nutrients), each cell in the population had an average doubling time of hours. To account for the limited availability of nutrients in the tumor microenvironment, we used a logistic growth model with a fixed carrying capacity. Cells in the population died at a fixed rate. Different EMP-associated phenotypes can exhibit different rates of cell division. Induction of EMT has been shown to arrest the cell cycle (Lovisa et al., 2015; Vega et al., 2004). Another study has shown that cells that have undergone a partial EMT can exhibit a hyper-proliferative phenotype (Handler et al., 2018). Here, we considered a simpler case wherein both division and death rates are independent of the phenotype of the cell. When a cell divides, the RNA and protein molecules in the parent cell are randomly partitioned among the daughter cells. Since *I*_*sig*_ represents multiple upstream signaling pathways whose functionality converges onto the regulatory circuit, the noise in the partitioning of *I*_*sig*_ among the daughter cells will be the dominant perturbation to the dynamics of the EMP regulatory circuit and is, therefore, the focus of our proposed model. The concentration of *I*_*sig*_ in each daughter cell after cell division was given by:

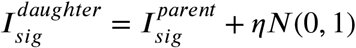

**Figure 1.**
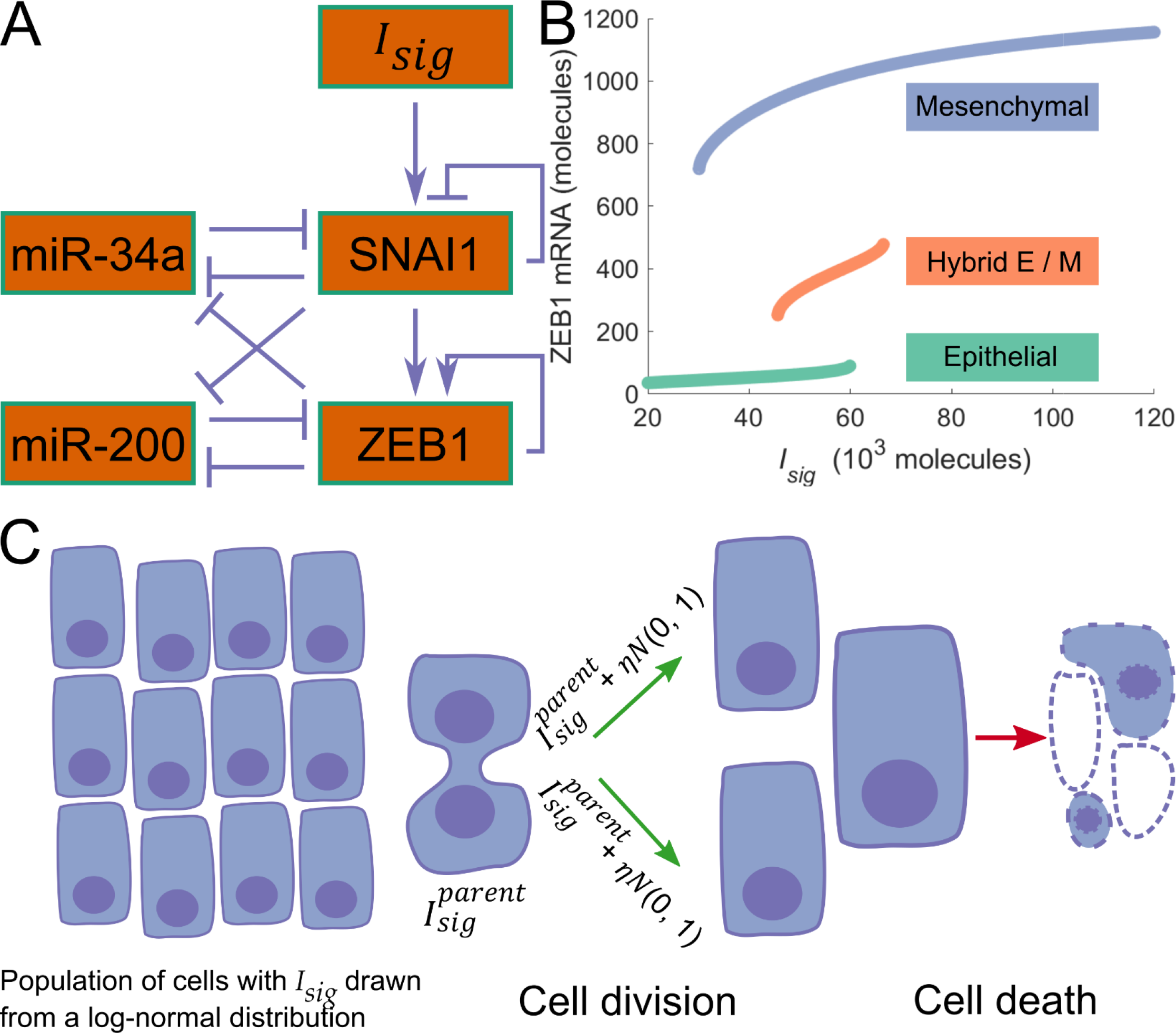
Description of the proposed model. **(A)** The regulatory circuit governing EMT and MET. **(B)** Bifurcation diagram of the EMT / MET regulatory circuit. **(C)** A schematic of the simulations carried out in the present study. Each cell carries a copy of the regulatory circuit shown in fig. 1 (A).

Here, *N*(0, 1) is the standard normal distribution and *η* is a model parameter controlling the variance of the noise distribution. The above equation models the typical scenario wherein division of the parent cell is preceded by gene duplication accompanied by near doubling of protein and RNA copy numbers. Each daughter cell then receives, on average, half the number of RNA and protein molecules present in the parent cell just before division. With the new concentration of *I*_*sig*_, the EMP regulatory circuit in the daughter cell may acquire a steady state different from that of the parent cell. The daughter cell can then exhibit a phenotype different from that of the parent cell before division. A schematic of the model is shown in fig. 1 (C).

### Cell-Phenotype Changes During EMP are Governed by the Noise in *I*_*sig*_ Partitioning During Cell Division

We started with a population of 500 cells on day 0. The initial concentration of *I*_*sig*_ in these cells was drawn from a log-normal distribution with median 2 × 10^4^ molecules / cell and with coefficient of variation 1.0. The dynamics of this population was simulated for a period of 8 weeks using the model of EMP described above, for different values of the noise parameter *η*. The cells grew quickly in number, with the total number of cells in the population becoming nearly stationary around day 18. The growth kinetics, as expected, were independent of the noise parameter *η* (fig. S1). Every time a cell divided, we recorded the phenotypes of the parent and the daughter cells. At low values of *η*, cells of all three phenotypes exhibited high rates of symmetric self-renewal i.e. both the daughter cells exhibited the same phenotype as the parent cell in most instances (fig. 2 (A)). As *η* was increased, the probability of a daughter cell acquiring a phenotype different from that of the parent cell increased for all three types of cells. For *η* = 1× 10^4^, there was a ~3% chance that a dividing epithelial cell will generate at least 1 daughter that is non-epithelial. The chances increased to ~22% for *η* = 3 × 10^4^. Similarly, for both hybrid E / M and mesenchymal cells, the probability of generating at least one daughter cell with a phenotype different from that of the parent cell increased significantly with an increase in *η*. Among the three phenotypes, hybrid E / M cells exhibited the highest probability of generating a daughter cell with a phenotype different from that of the parent cell. Even at low values of *η*, the probability that at least one of the daughter cells will be non-hybrid E / M when a hybrid E / M cell divided was as high as 50%. This probability increased further with an increase in *η*, reaching ~82% for *η*=3×10^4^. On the other hand, mesenchymal cells exhibited the least probability of generating a daughter cell with a phenotype different from that of the parent cell. The probability of generating a non-mesenchymal cell during mesenchymal cell division saturated around 10% with an increase in *η*.

The model can predict how the fractions of different phenotypes in a population of cells will evolve over time, given the initial phenotypic composition of the population. Examples of temporal dynamics for different values of *η* are shown in fig. 2 (B). On starting with a population that is ~84% epithelial, the fraction of epithelial cells in the population declined over time. The rate of this decline increased with increasing *η* due to the increase in the probability of generating a non-epithelial daughter cell during epithelial cell division. The fraction of mesenchymal cells, on the other hand, increased with time as mesenchymal cells were generated via the division of epithelial cells. Given the high probability of symmetric self-renewal of mesenchymal cells even at high values of *η*, the fraction of these cells in the population increased with increasing *η*. The temporal dynamics of the hybrid E / M fraction was non-monotonic and, due to the high probability of generating a non-hybrid E / M cell during the division of a hybrid E / M cell, their fraction in the population remained below 10%.

**Figure 2.**
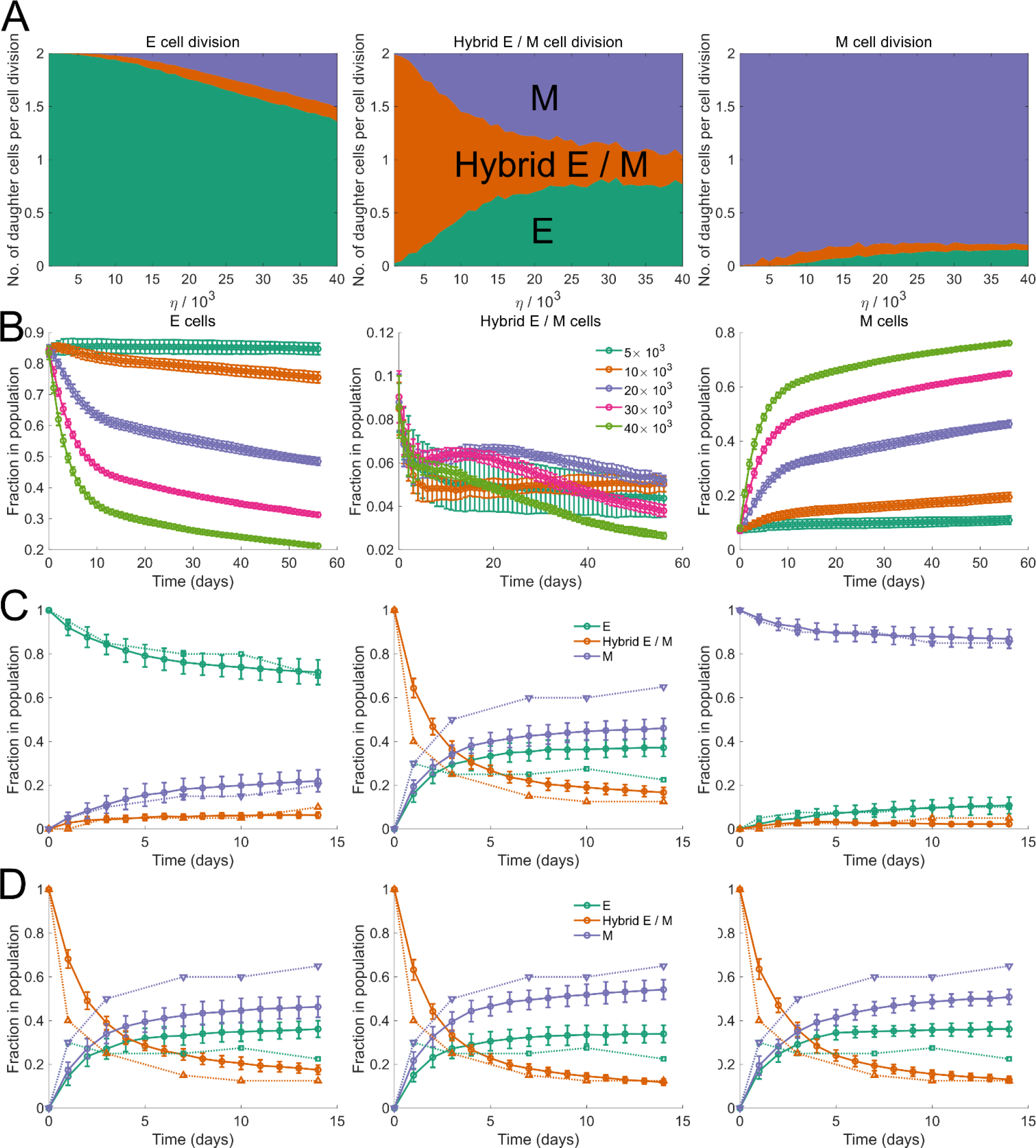
Dynamics of the model. **(A)** Average number of daughter cells of each phenotype generated per cell division when the parent cell was an epithelial cell, a hybrid E / M cell, or a mesenchymal cell. **(B)** Temporal dynamics of the fraction of epithelial (E) cells, hybrid E / M cell, and mesenchymal (M) cells in a population of cancer cells. Different colors indicate different values of the noise parameter *η*. Also, see fig. S1. **(C)** Fractions of different EMP-associated phenotypes assessed at different time points during a two-week period. Dashed curves, with inverted triangles, represent the fractions of different phenotypes re-plotted from Ruscetti *et al.* (Ruscetti et al., 2016). Solid curves, with circles, represent the predictions from our model using *η* = 1.9 × 10^4^. **(D)** Fractions of EMP-associated phenotypes at different time points in a population of cancer cells which consisted of only hybrid E / M cells on day 0 under three regimes: asymmetric distribution of miR-34a among the daughter cells during the division of an epithelial cell (left panel; *η* = 2.1 × 10^4^), asymmetric distribution of miR-34a among the daughter cells during the division of a hybrid E / M cell (center panel; *η* = 1.9 × 10^4^), and asymmetric distribution of miR-34a among the daughter cells during the division of a mesenchymal cell (right panel; *η* = 2.0 × 10^4^). Also, see fig. S2 and S3. In each panel, error bars indicate the standard deviation calculated over 16 independent runs.

### Model Captures the Temporal Changes in Phenotypic Composition in a Mouse Model of Prostate Cancer

We tested the predictions of our model in tumor cells isolated from murine prostate tumors (Ruscetti et al., 2016). To study the role of EMP in the distant metastasis of prostate cancer *in vivo*, Ruscetti *et al.* crossed *Pb-Cre*^+/−^; *Pten*^*L/L*^; *Kras*^*G12D*/+^ mice with *Vim-GFP* reporter mice to track EMP in prostate cancer cells. EpCAM^+^GFP^−^ epithelial tumor cells were FACS (fluorescence-activated cell sorting) sorted from 10-week old prostates and cultured *in vivo* (*PKV* cell line). FACS sorting of these cells after 14 days of *in vitro* culture revealed three distinct phenotypes-EpCAM^+^GFP^−^ (epithelial), EpCAM^−^GFP^+^ (mesenchymal), and EpCAM^+^GFP^+^ (hybrid E / M). Epithelial, mesenchymal, and hybrid E / M cells FACS sorted from the *PKV* cell line were then cultured *in vitro*. Fractions of different phenotypes in each culture at different time points were tracked using FACS. The temporal dynamics has been replotted in fig. 2 (C) along with the predictions from our model.

Model predictions shown in fig. 2 (C) were obtained by varying the noise parameter *η* to minimize the root mean square deviation from experimental data. Our model was able to capture the time scale over which the phenotypic composition of the population changed in each of the three cases with different initial compositions. When starting with a population consisting of only epithelial cells, ~70% of cells were epithelial at the end of the 14-day simulation and experiment period. Consistent with the high rate of symmetric self-renewal of mesenchymal cells, nearly 87% of the population was still mesenchymal after the 14-day period when starting with a population of only mesenchymal cells on day 0. On the other hand, on starting with a population of hybrid E / M cells on day 0, the fraction of these cells in the population dropped below 40% within 3 days. A mixture of epithelial and mesenchymal cells had quickly been generated by the population of hybrid E / M cells.

Hybrid E / M cells isolated from the *PKV* cell line, when cultured *in vitro*, generated a population with more mesenchymal cells than epithelial cells (fig. 2 (C); center panel). This behavior was not captured by our model which treats epithelial, mesenchymal, and hybrid E / M cells equivalently i.e. doubling times, death rates, and the characteristic of noise in the partitioning of *I*_*sig*_ during cell division are identical for cells of all three phenotypes. Hybrid E / M cells, however, exhibit a special property— multiple studies have reported that the hybrid E / M phenotype is associated with the expression of cancer stem cell markers (Bocci et al., 2019; Grosse-Wilde et al., 2015; Jordan et al., 2011; Schmidt et al., 2015). During the division of colon cancer stem cells (Bu et al., 2013, 2016; Wang et al., 2016) and of mammary stem cells (Bonetti et al., 2019), asymmetric partitioning of miR-34a among the daughter cells has been observed.

We incorporated this property of hybrid E / M cells into our model by setting the concentration of miR-34a in one of the daughter cells to zero during the division of a hybrid E / M cell while keeping the concentration of miR-34a in the other daughter cell same as the concentration in the parent cell. When simulating the dynamics of a population with only hybrid E / M cells on day 0, this modification to the model led to a larger fraction of mesenchymal cells as compared to epithelial cells in the population at subsequent time points. This modification allowed for a better fit to the experimental observations obtained for cells isolated from the *PKV* cell line (fig. 2 (D); center panel and fig. S2). Asymmetric division of miR-34a among the daughter cells during the division of epithelial or mesenchymal cells did not typically lead to a better fit to experimental data (fig. 2 (D); left and right panels and fig. S2).

This modified model, with asymmetric division of miR-34a among the daughter cells during the division of a hybrid E / M cell, has been used to obtain all subsequent results reported in this paper.

### Hysteresis in the Dynamics of Epithelial-Mesenchymal Plasticity

The regulatory circuit driving EMP, shown in fig. 1 (A), exhibits nonlinear dynamics. Such systems can exhibit hysteresis wherein the system response depends not only on the present input but also on the input history. We investigated if hysteresis can also arise in the dynamics of fractions of different phenotypes in a population of cancer cells. Starting with a population of epithelial cells on day 0, a fixed dosage of *I*_*sig*_ was added each day for 10 days. A fixed dosage of *I*_*sig*_ was then withdrawn over the next 10 days. Fig. 3 shows the fractions of different phenotypes in the population over the 20 day period. Since *I*_*sig*_ promotes EMT, the fraction of mesenchymal cells in the population increased over the first 10 days from 0% to >96% while the fraction of epithelial cells declined to less than 3%. When *I*_*sig*_ was withdrawn in fixed dosages day 11 onwards, this trend was reversed. However, the fractions of epithelial and mesenchymal cells in the population from day 11 to day 20 did not retrace the behavior recorded from day 0 to day 10. On day 20, ~40% of the cells still exhibited a mesenchymal phenotype which was maintained in the absence of additional *I*_*sig*_ dosages. These cells thus represented a population that underwent an EMT which was irreversible on the time scale considered here (fig. S4). Hysteretic control of EMT and MET dynamics was recently confirmed in multiple normal and cancerous mammary epithelial cell lines including MCF10A, MCF7, HMLE, T47D, and 4T1 (Celià-Terrassa et al., 2018). The study also confirmed the persistence of the mesenchymal phenotype long after the withdrawal of the EMT-inducing signal as predicted by our model (fig. S4).

**Figure 3.**
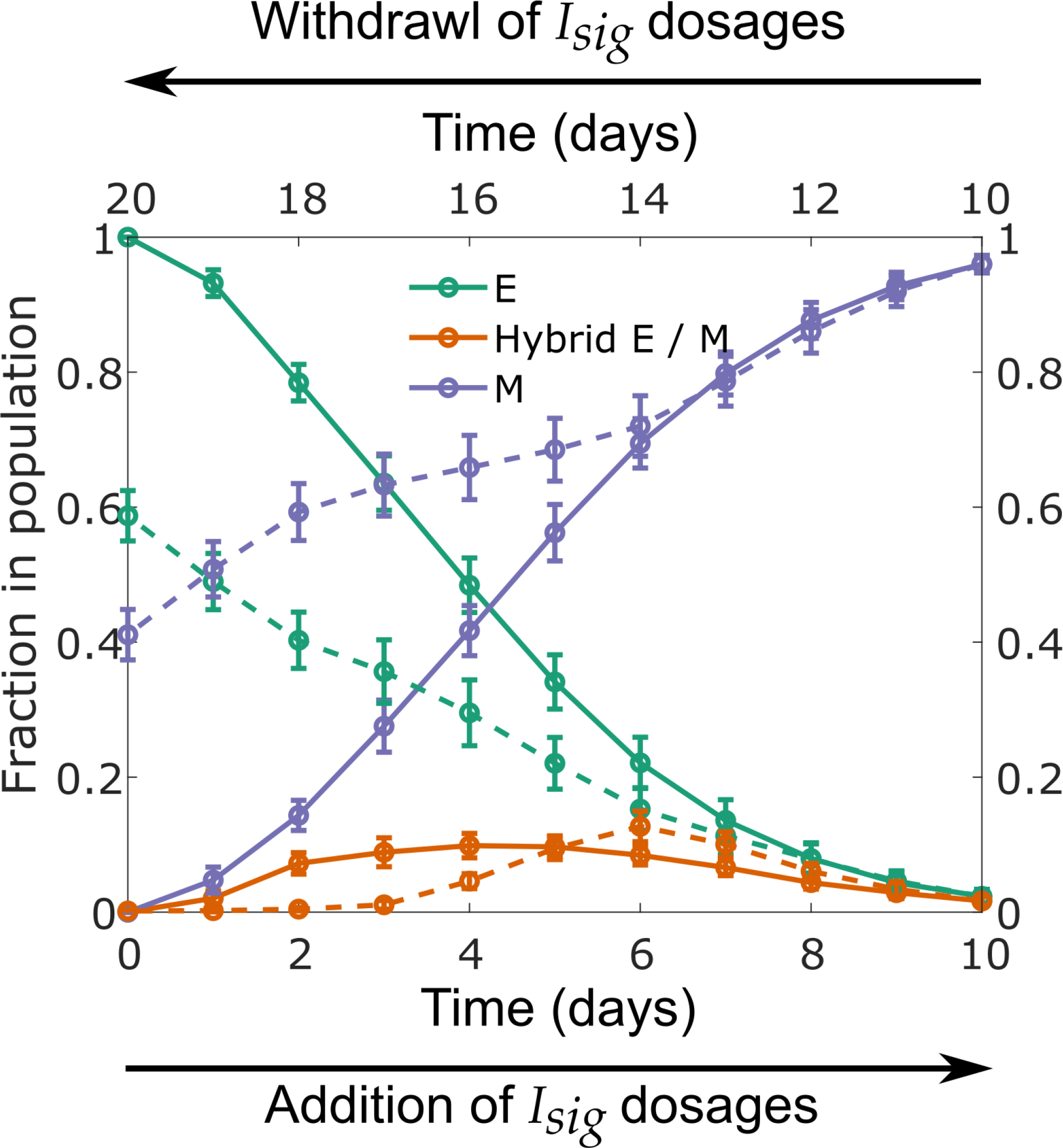
Hysteresis in the temporal behavior of fractions of different phenotypes. Solid curves show the behavior from day 0 to day 10 (bottom horizontal axis) and the dashed curves show the behavior from day 11 to day 20 (top horizontal axis). Here, *η* = 1.9 × 10^4^. Also, see fig. S4 and fig. S5.

### Frequencies of Spontaneous Cell-Phenotype Changes can be Regulated by Modifying the Regulatory Circuit Driving EMP

In our model, cell-phenotype changes occur, in response to noise, due to the tri-stability in the dynamics of the EMP regulatory circuit. Thus, modifications to this regulatory circuit which change the bifurcation diagram (fig. 1(B)) should alter the frequencies of cell-phenotype changes. Mathematical models have shown that the range of *I*_*sig*_ levels for which a stable hybrid E / M phenotype can exist is much larger when GRHL2 is coupled with the EMP regulatory circuit (Jolly et al., 2016) or when ΔNP63α is added to the regulatory circuit (Jolly et al., 2017) (fig. 4(A)). In both cases, given the larger range of *I*_*sig*_ levels for which a stable hybrid E / M phenotype can exist, we expected the probability of generating an epithelial or mesenchymal daughter cell during the division of a hybrid E / M cell to decrease upon the inclusion of GRHL2 or ΔNP63α in our model. This was confirmed by tracking the phenotypes of daughter cells formed via the division of a hybrid E / M cell upon including the activity of GRHL2 or ΔNP63α. For all values of the noise parameter *η* investigated, a higher probability of generating at least 1 daughter hybrid E / M cell during the division of a hybrid E / M cell was observed upon the inclusion of both GRHL2 and ΔNP63α (fig. 4 (B)). Even at high values of *η*(*η*=4 ×10^4^),inclusion of ΔNP63α did not let the probability of generating at least 1 non-hybrid E / M daughter cell exceed 39%. This probability was > 86% in the absence of ΔNP63α. Upon starting with a population of hybrid E / M cells on day 0,nearly 30% of cells exhibited a hybrid E / M phenotype on day 14 upon the inclusion of GRHL2 as compared to less than 12% of cells in the absence of GRHL2 activity (fig. 4(C);left panel). When ΔNP63α was included, the fraction of hybrid E / M cells in the population on day 14 was ~56%(fig. 4 (C); right panel), a fraction larger than the one seen in the presence of GRHL2 activity or in the absence of both GRHL2 and ΔNP63α. This behavior is reminiscent of the increased mean residence time of cells in a hybrid E / M state in the presence of GRHL2 and ΔNP63α which has been reported previously (Biswas et al., 2019).

**Figure 4.**
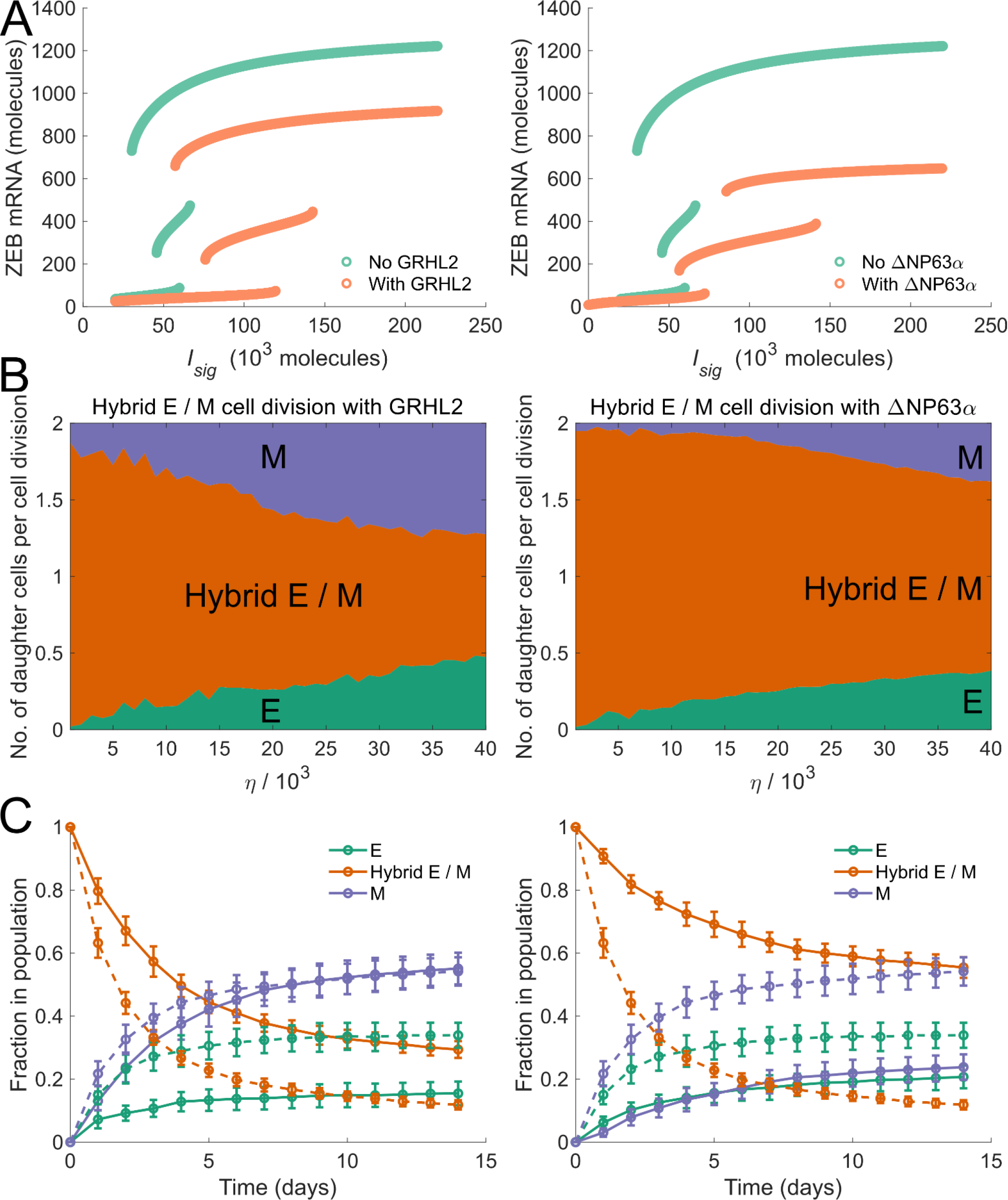
Model dynamics in the presence of EMT modulators. **(A)** Bifurcation diagrams of the regulatory circuit governing EMT / MET in the presence of GRHL2 (left panel) and in the presence of ΔNP63*α* (right panel). **(B)** Average number of daughter cells of each phenotype generated per cell division during the division of a hybrid E / M cell in the presence of GRHL2 (left panel) and in the presence of ΔNP63*α* (right panel). **(C)** Fractions of EMP-associated phenotypes at different time points in a population of cancer cells which consisted of only hybrid E / M cells on day 0— dynamics in the presence of GRHL2 (solid curves, left panel) and dynamics in the presence of ΔNP63*α* (solid curves, right panel). Dashed curves in left and right panels indicate the fractions of different phenotypes in the absence of GRHL2 and ΔNP63*α* activity respectively. Here, *η* = 1.9 × 10^4^. In each panel, error bars indicate the standard deviation calculated over 16 independent runs.

We next investigated how other EMP modulators may alter the phenotypic composition of a population of cells. Biddle *et al.* used co-treatment with 5*μ*M retinoic acid and 0.5ng / ml TGF-*β* to maintain a subpopulation of cells in the partial EMT state for two oral squamous cell carcinoma cell lines (Biddle et al., 2016). We used our model to characterize the effects of co-treatment with retinoic acid, an MET inducer (Wu et al., 2017), and exogenous TGF-*β*, an EMT inducer (Nieto et al., 2016). Starting with a population of epithelial cells on day 0, we simulated the model for different concentrations of retinoic acid and exogenous TGF14-*β*. Fig. 5 (A) shows the fractions of epithelial, mesenchymal, and hybrid E / M cells in the population on day. The model predicts that using a certain combinations of retinoic acid and TGF-*β* concentrations, a population with a large fraction of cells that have undergone a partial EMT can be maintained, thereby recapitulating the experimental observations of Biddle *et al.* (Biddle et al., 2016) (fig. 5 (B)).

**Figure 5.**
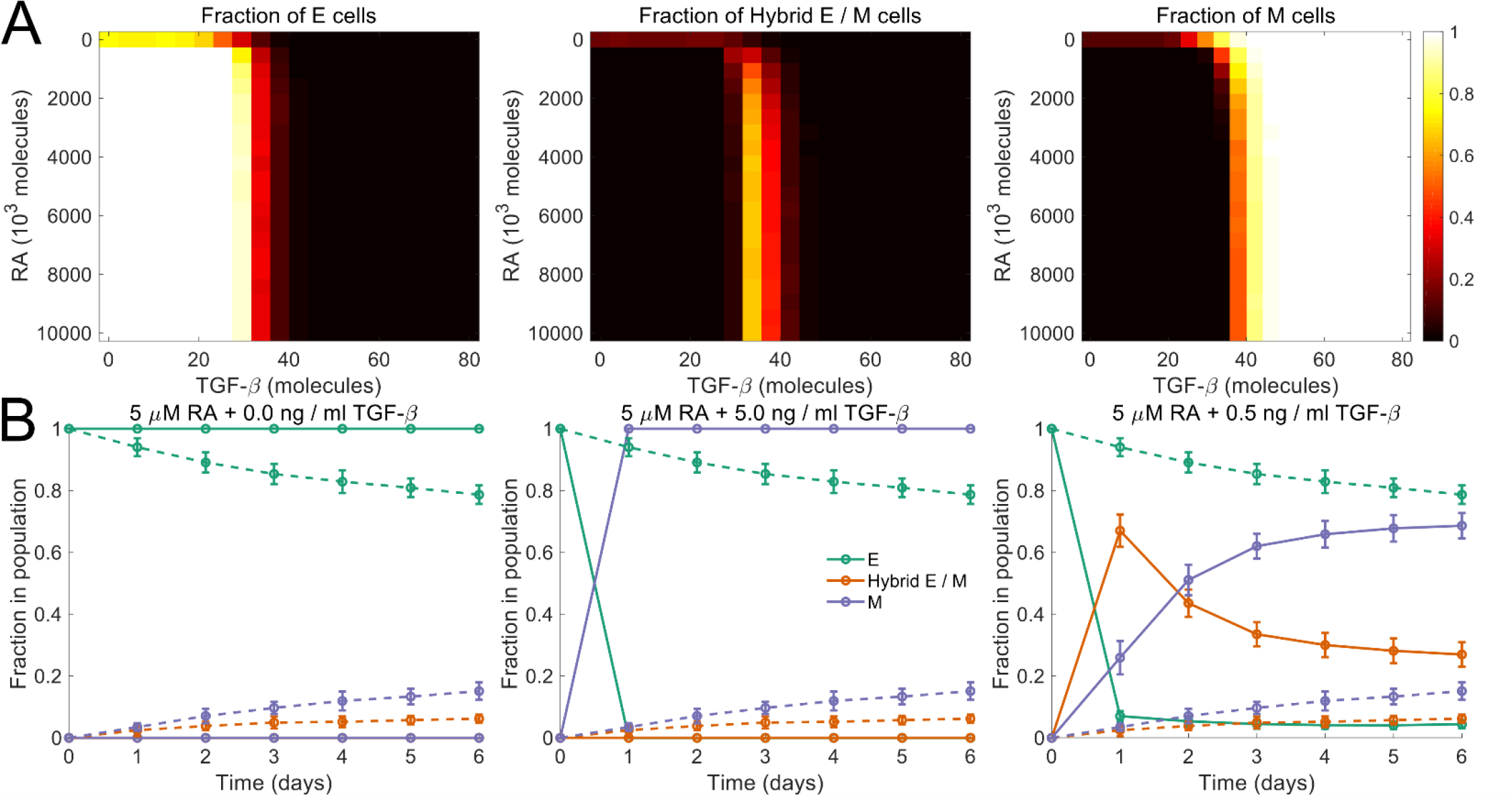
Model dynamics in the presence of retinoic acid and TGF-*β*. **(A)** Fractions of different phenotypes in a population after 14 days upon treatment with different concentrations of retinoic acid (RA) and TGF-*β*. The population consisted of only epithelial cells on day 0. **(B)** Biddle *et al.* characterized the plasticity of oral squamous cell carcinoma cells in response to co-treatment with different concentrations of retinoic acid (RA) and TGF-*β*. 5.0 μM RA blocked EMT. Its effect was completely abrogated by the addition of 5.0 ng / ml of TGF-*β*. Co-treatment with 5 μM RA and 0.5 ng / ml TGF-*β* established a regime in which a population of cells that have undergone a partial EMT was maintained. Starting with a population of epithelial cells on day 0, we simulated the dynamics under these three co-treatment regimens. The predictions from our model are in agreement with the experimental observations of Biddle *et al.* (Biddle et al., 2016). Dashed curves indicate the dynamics in the absence of both RA and TGF-*β*. In each panel, error bars indicate the standard deviation calculated over 16 independent runs. Here, *η* = 1.9 × 10^4^.

### Drugs Targeting Epithelial and Mesenchymal Cells Have a Synergistic Effect in Restricting Tumor Size

Finally, we investigated the effectiveness of targeting different phenotypes or phenotype combinations in restricting tumor growth. Drug activity was modeled using the shifted Hill function which altered the death rate of cancer cells by a multiplicative factor. Fig. 6 shows how the number of tumor cells in the population changed under different drug regimens. Drugs targeting a single phenotype, whether epithelial, mesenchymal, or hybrid E / M, had a negligible effect on tumor size, even at high concentrations. Since epithelial cells can generate a population of mesenchymal cells, targeting the epithelial phenotype led to an increase in the fraction of mesenchymal cells in the population. These cells exhibit a low probability of generating non-mesenchymal daughter cells and had greater nutrient availability once the population of epithelial cells had been depleted by the drug. Similarly, targeting of mesenchymal cells increased the fraction of epithelial cells in the population while doing little to restrict the tumor size. Targeting the hybrid E / M phenotype, too, had little effect due to the high probability of generating epithelial or mesenchymal daughter cells during hybrid E / M cell division. This behavior would allow for the rapid generation of epithelial and mesenchymal cells in the population, both of which are not targeted by the drug. Co-targeting of epithelial and hybrid E / M cells or that of mesenchymal and hybrid E / M cells did not substantially alter the tumor size either. The un-targeted phenotype took over the population in both cases.

**Figure 6.**
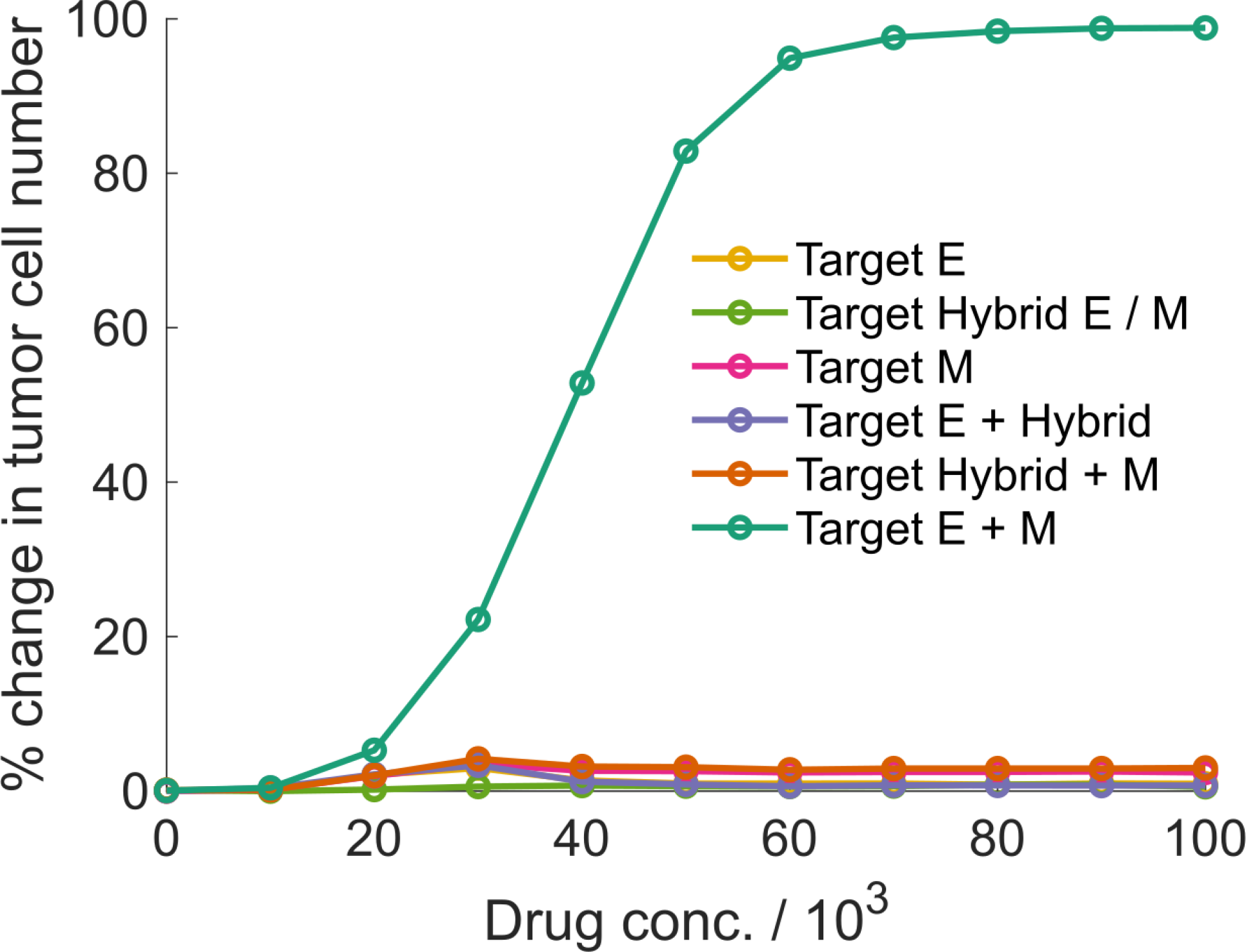
Percentage change in the number of tumor cells after drug treatment for 28 days. Different colors indicate different treatment regimens.

A drastic decrease in tumor size was observed upon co-targeting of epithelial and mesenchymal cells (fig. 6). While the resource availability for hybrid E / M cells went up in this regime due to the depletion of epithelial and mesenchymal populations, hybrid E / M cells, with low rates of symmetric self-renewal, quickly generated a population with epithelial and mesenchymal cells. Both these phenotypes were subsequently targeted. This led to a drastic decline in tumor size and the decline increased with increase in drug dosages. Under this regime, the phenotypic composition of the tumor cell population was also fundamentally altered— fractions of the three phenotypes in the population become nearly equal (fig. 7). Thus, simultaneous targeting of epithelial and mesenchymal cells, while effective in restricting tumor growth, can bolster phenotypic heterogeneity and increase the fraction of hybrid E / M cells in the population. This phenotype has been implicated in multiple metastatically aggressive behaviors (Jolly et al., 2015). Thus, the treatment regime, though effective in restricting tumor growth, will have significant implications for cancer relapse post-treatment.

**Figure 7.**
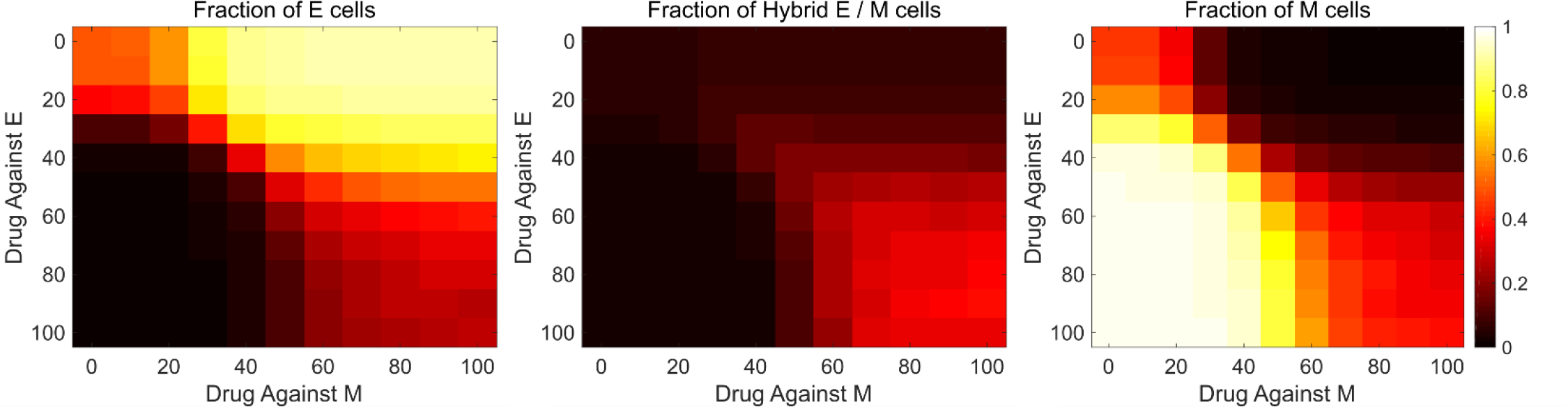
Change in phenotypic composition of the tumor upon drug treatment. Fractions of different phenotypes in a population of cancer cells upon co-treatment, for 28 days, with varying concentrations of drugs targeting epithelial and mesenchymal cells. On day 0, all cells in the population were epithelial. Drug concentrations are indicated as 10^3^ units.

### Discussion

Multiple studies have characterized the phenotypic heterogeneity in cancer cells (Meacham and Morrison, 2013). However, the ability of cancer cell populations to retain this heterogeneity over extended periods of time and through multiple generations and passages has remained a mystifying feature of the disease with wide-ranging implications for the design of anti-cancer therapeutic strategies. Here, we have described a model for the generation and maintenance of epithelial-mesenchymal heterogeneity in a population of cancer cells. Our model is based upon the tri-stability of the regulatory circuit driving EMT and MET, and thus connects the dynamical behavior of the regulatory mechanism within each cell to the phenotypic composition of the population. While the model does not capture the rich behavior that may arise from the variable strengths of regulatory interactions in cells in a population or from cell-cell communication, it is a useful first step in understanding the generation and maintenance of phenotypic heterogeneity in cancer cells. Targeting the ability of cancer cells to change phenotypes and generate heterogeneous populations has recently been proposed as a therapeutic strategy for combating drug resistance (Risom et al., 2018). Our model can be a platform to identify putative therapeutic targets for inhibiting the generation, maintenance, and propagation of phenotypic heterogeneity in cancer cells that exhibit EMP.

A key prediction of our model is that noise in the partitioning of parent cell biomolecules among the daughter cells at the time of cell division is sufficient for the generation of phenotypic heterogeneity in a population of cancer cells. Since the heterogeneity arises from cell division-associated noise, it can be maintained over multiple generations and can be propagated from a small population. This makes it possible for a small population of tumor cells, like that leftover after surgery or a chemotherapeutic regime, to recreate the full-fledged phenotypic heterogeneity of a mature tumor if conditions that can sustain tumor growth emerge at a later point in time. The hybrid E / M phenotype can quickly generate a mixed population of epithelial and mesenchymal cells. This behavior may contribute to the metastatic aggressiveness widely associated with the hybrid E /M phenotype (Jolly et al., 2015) and may account for the abundance of these cells in aggressive disease subtypes such as inflammatory breast cancer (Jolly et al., 2017). Hybrid E / M cells, when disseminated to a distant organ, can quickly generate epithelial cells to colonize the tissue. Further, these cells can potentially re-create the phenotypic complexity of the primary tumor at the distant organ site, providing a suitable microenvironment for tumor growth in the new tissue. Our model thus highlights how the initial phenotypic composition can affect the temporal population-level behavior of tumor cell populations and the implications of such behavior for cancer metastasis.

Another key prediction of our model is hysteresis in the temporal dynamics of fractions of different phenotypes in a population of cancer cells. Note that the hysteretic behavior reported here is distinct from the behavior observed in typical non-linear systems where the steady state reached by a system can depend not only on the final system input but also on the input history. In our model, the population of cells does not reach a steady state of fractions of different phenotypes within the time period over which most experimental studies are carried out. Hysteresis is rather observed in the trajectories displayed by the fractions of different phenotypes over experimental time scales, a behavior that has been observed in multiple cell lines (Celià-Terrassa et al., 2018). Our model further predicts that some cancer cells undergo a seemingly irreversible EMT upon transient exposure to an EMT promoter such as TGF-*β*. These cells can retain a mesenchymal phenotype long after the EMT-inducing signal has been withdrawn and do not revert back to an epithelial state during the time period considered in the present study. Such a time period is typical of most experimental studies. An irreversible EMT upon chronic exposure to EMT-inducing signals has been reported (Katsuno et al., 2019; Zhang et al., 2014). Our model shows that even transient activation of EMT signaling in cancer cells can generate heterogeneity which will be maintained and propagated long after the exogenous signal has been withdrawn. Such behavior has been confirmed in a recent study which showed that transient exposure to TGF-*β* can be sufficient to generate cells with a mesenchymal phenotype and such cells retain the acquired mesenchymal phenotype for days after TGF-*β* withdrawal (Celià-Terrassa et al., 2018).

Finally, our model provides significant insight into how phenotypically heterogeneous populations will respond to drugs. When cells that can generate cells of other phenotypes are treated with a drug that targets only one phenotype, untargeted cells, generated from the drug-sensitive cells, will take over the population with no significant decline in tumor size. Similar behavior has been described in the context of adaptive cancer therapy wherein the small number of chemotherapy-resistant cells present in the tumor can take over the population once the chemo-sensitive population has been killed off (Gatenby et al., 2009). In our model, under drug activity, cells can switch between drug-sensitive and drug-resistant phenotypes which markedly alters the population response to the drug. Leaving one of the more stable phenotypes untargeted will lead to that phenotype taking over the population with no decline in tumor size. In the context of EMP, this is observed in the scenario wherein either epithelial or mesenchymal cells are left untargeted. Co-targeting both these phenotypes resulted in the best therapeutic response with respect to decrease in tumor size. However, even in this regime, a small, highly heterogeneous population is left behind which can quickly regenerate the full-fledged phenotypic heterogeneity of a mature tumor once the drug treatment has been withdrawn. Thus, our model highlights the difficulty in developing effective therapeutic strategies for the heterogenous disease that is cancer.

While the present study focuses on epithelial-mesenchymal heterogeneity in cancer cells, the model can easily be generalized to describe spontaneous changes in the phenotypic composition of a cancer cell population in multiple scenarios wherein the regulatory mechanism driving different phenotypes has been characterized. Our model can easily be adapted to describe stem-cell heterogeneity in the tumor microenvironment (Bocci et al., 2019), neuroendocrine plasticity in small cell lung cancer (Wooten et al., 2018), and luminal-basal plasticity in triple-negative breast cancer (Risom et al., 2018). Further, the model can be extended to describe spatio-temporal heterogeneity generated via a juxtacrine signaling mechanism such as Notch signaling in small cell lung cancer (Lim et al., 2017).

## Methods

### Computer simulation of the model

The dynamics of the regulatory circuit was simulated using ordinary differential equations (ODEs). The mathematical form of the ODEs and the relevant kinetic parameters are described in the supplementary text. Population dynamics were simulated using Gillespie’s algorithm (Gillespie, 1977). The model parameters are listed in the supplementary text. C++ code used to generate the figures presented in the study is available online on GitHub (https://github.com/st35/cancer-EMT-heterogeneity-noise).

## Acknowledgements

This work was supported by the National Science Foundation grants PHY-1427654 and PHY-1605817. M.K.J. is supported by the Ramanujan Fellowship, awarded by the Science and Engineering Research Board, Department of Science and Technology, Government of India (SB/S2/RJN-049/2018).

## Author Contributions

Conceptualization, S.T. and M.K.J.; Methodology, S.T. and M.K.J.; Investigation, S.T.; Writing-Original Draft, S.T.; Writing-Review & Editing, S.T., H.L., and M.K.J.; Supervision, H.L. and M.K.J.

## Declaration of Interests

The authors declare no competing interests.

