## Supplementary Information for "A Mechanism for Epithelial-Mesenchymal Heterogeneity in a Population of Cancer Cells"

### Dynamics of the EMT / MET regulatory circuit

The regulatory circuit driving EMT and MET shown in fig. 1 (A) consists of two interconnected modules. The first module involves mutual inhibitory interactions between miR-34a and *SNAI1*. In the second module, miR-200 and *ZEB1* form a mutual inhibitory loop. The circuit thus includes both transcriptional and post-transcriptional regulation. The dynamics of this circuit has been described in detail previously (Lu et al., 2013). We used the same approach to model the behavior of the regulatory circuit. The ordinary differential equations (ODEs) that describe the dynamics of this regulatory circuit along with the relevant kinetic parameters were obtained from Lu *et al.* (Lu et al., 2013). The ODEs are listed below.

$$\frac{d\mu_{200}}{dt}=g_{\mu_{200}}H\left( Z, \lambda_{Z}^{\mu_{200}} \right)H\left( S, \lambda_{S}^{\mu_{200}} \right)-m_{Z}Y_{\mu}\left( \mu_{200} \right)-k_{\mu_{200}}\mu_{200}$$

$$\frac{dm_{Z}}{dt}=g_{m_{Z}}H\left( Z, \lambda_{Z}^{m_{Z}} \right)H\left( S, \lambda_{S}^{m_{Z}} \right)-m_{Z}Y_{m}\left( \mu_{200} \right)-k_{m_{Z}}m_{Z}$$

$$\frac{dZ}{dt}=g_{Z}m_{Z}L\left( \mu_{200} \right)-k_{Z}Z$$

$$\frac{d\mu_{34}}{dt}=g_{\mu_{34}}H\left( S,\lambda_{S}^{\mu_{34}} \right)H\left( Z, \lambda_{Z}^{\mu_{34}} \right)-m_{S}Y_{\mu}\left( \mu_{34} \right)-k_{\mu_{34}}\mu_{34}$$

$$\frac{dm_{S}}{dt}=g_{m_{S}}H\left( S, \lambda_{S}^{m_{S}} \right)H\left( I_{sig}, \lambda_{I_{sig}}^{m_{S}} \right)-m_{S}Y_{m}\left( \mu_{34} \right)-k_{m_{S}}m_{S}$$

$$\frac{dS}{dt}=g_{S}m_{S}L\left( \mu_{34} \right)-k_{S}S$$

$$\frac{dI}{dt}=0.0$$

Here, $\mu_{200}=$ [miR-200], $m_{Z}=$ [ZEB1 mRNA], $Z=$ [ZEB1], $\mu_{34}=$ [miR-34a], $m_{S}=$ [SNAI1 mRNA], $S=$ [SNAI1], and $I=$ [$I_{sig}$]. [$\cdot$] represents the concentration of a molecular species within a cell. $H$ is the shifted Hill function.

$$H(B, \lambda)=\lambda+\frac{1.0-\lambda}{1.0+\left( \frac{B}{B_{0}} \right)^{n_{B}}}$$

The functions $Y_{\mu}$, $Y_{m}$, and $L$ describe the post-transcriptional regulation of mRNA activity by micro-RNAs and have been described previously (Lu et al., 2013).

$$L\left( \mu\right)=\sum_{i=0}^{n} \binom{n}{i}l_{i}M_{n}^{i}(\mu)$$

$$Y_{m}\left( \mu\right)=\sum_{i=0}^{n} \binom{n}{i}\gamma_{m_{i}}M_{n}^{i}(\mu)$$

$$Y_{\mu}\left( \mu\right)=\sum_{i=0}^{n} \binom{n}{i}\gamma_{\mu_{i}}M_{n}^{i}(\mu)$$

$$M_{n}^{i}\left( \mu\right)=\frac{\left( \frac{\mu}{\mu^{0}} \right)^{i}}{\left( 1.0+\frac{\mu}{\mu^{0}} \right)^{n}}$$

Here, $\mu$ is the concentration of the micro-RNA and $n$ is the number of micro-RNA binding sites on the mRNA. For the inhibition of *SNAI1* mRNA by miR-34a, $n=2$. For the inhibition of *ZEB1* mRNA by miR-200, $n=6$. The values of all kinetic parameters are listed in tables 1 and 2.

### Dynamics in the presence of EMT modulators

Fig. SI 1 shows how different EMT modulators couple with the regulatory circuit shown in fig. 1 (A). The studies that describe the activity of these EMT modulators are cited in the caption of fig. SI 1. ODEs listed in the previous section were modified to incorporate the activity of the respective EMT modulators.

**GRHL2**

$$\frac{dm_{Z}}{dt}=g_{m_{Z}}H(G, \lambda_{G}^{m_{Z}})H\left( Z, \lambda_{Z}^{m_{Z}} \right)H\left( S, \lambda_{S}^{m_{Z}} \right)-m_{Z}Y_{m}\left( \mu_{200} \right)-k_{m_{Z}}m_{Z}$$

$$\frac{dm_{G}}{dt}=g_{m_{G}}H\left( Z,\lambda_{Z}^{m_{G}} \right)-k_{m_{G}}m_{G}$$

$$\frac{dG}{dt}=g_{G}m_{G}-k_{G}G$$

Here, $G=$ [GRHL2] and $m_{G}=$ [GRHL2 mRNA].

$\boldsymbol{\Delta}$**NP63**$\boldsymbol{\alpha}$

$$\frac{dm_{Z}}{dt}=g_{m_{Z}}H(\mu_{205}, \lambda_{\mu_{205}}^{m_{Z}})H\left( Z, \lambda_{Z}^{m_{Z}} \right)H\left( S, \lambda_{S}^{m_{Z}} \right)-m_{Z}Y_{m}\left( \mu_{200} \right)-k_{m_{Z}}m_{Z}$$

$$\frac{dm_{S}}{dt}=g_{m_{S}}H(P_{63}, \lambda_{P_{63}}^{m_{S}})H\left( S, \lambda_{S}^{m_{S}} \right)H\left( I_{sig}, \lambda_{I_{sig}}^{m_{S}} \right)-m_{S}Y_{m}\left( \mu_{34} \right)-k_{m_{S}}m_{S}$$

$$\frac{d\mu_{205}}{dt}=g_{\mu_{205}}H\left( P_{63}, \lambda_{P_{63}}^{\mu_{205}} \right)-k_{\mu_{205}}\mu_{205}$$

$$\frac{dP_{63}}{dt}=0.0$$

Here, $\mu_{205}=$ [miR-205] and $P_{63}=$ [$\Delta$NP63$\alpha$]

**Retinoic acid and TGF-**$\boldsymbol{\beta}$

$$\frac{dm_{Z}}{dt}=g_{m_{Z}}H(T, \lambda_{T}^{m_{Z}})H\left( Z, \lambda_{Z}^{m_{Z}} \right)H\left( S, \lambda_{S}^{m_{Z}} \right)-m_{Z}Y_{m}\left( \mu_{200} \right)-k_{m_{Z}}m_{Z}$$

$$\frac{d\mu_{200}}{dt}=g_{\mu_{200}}H(R, \lambda_{R}^{\mu_{200}})H\left( Z, \lambda_{Z}^{\mu_{200}} \right)H\left( S, \lambda_{S}^{\mu_{200}} \right)-m_{Z}Y_{\mu}\left( \mu_{200} \right)-k_{\mu_{200}}\mu_{200}$$

$$\frac{dR}{dt}=0.0$$

$$\frac{dT}{dt}=0.0$$

Here, $R=$ [RA] and $T=$ [TGF-$\beta$].

### Simulation of population dynamics

The dynamics of the population of cancer cells was simulated using Gillespie’s algorithm (Gillespie, 1977). The dynamics involved two types of events: cell division and cell death. Following logistic growth, the rate of cell division at time $t$ was given as

$$r(t)=r_{0}(1-\frac{N(t)}{K})$$

where $r_{0}=0.0182$ per hour is the rate of cell division in the absence of any competition (or, equivalently, in the presence of an infinite supply of nutrients), $N(t)$ is the total number of cells in the population at time $t$, and $K$ is the total carrying capacity.

The death rate was $0.00182$ per hour (one-tenth the value of $r_{0}$).

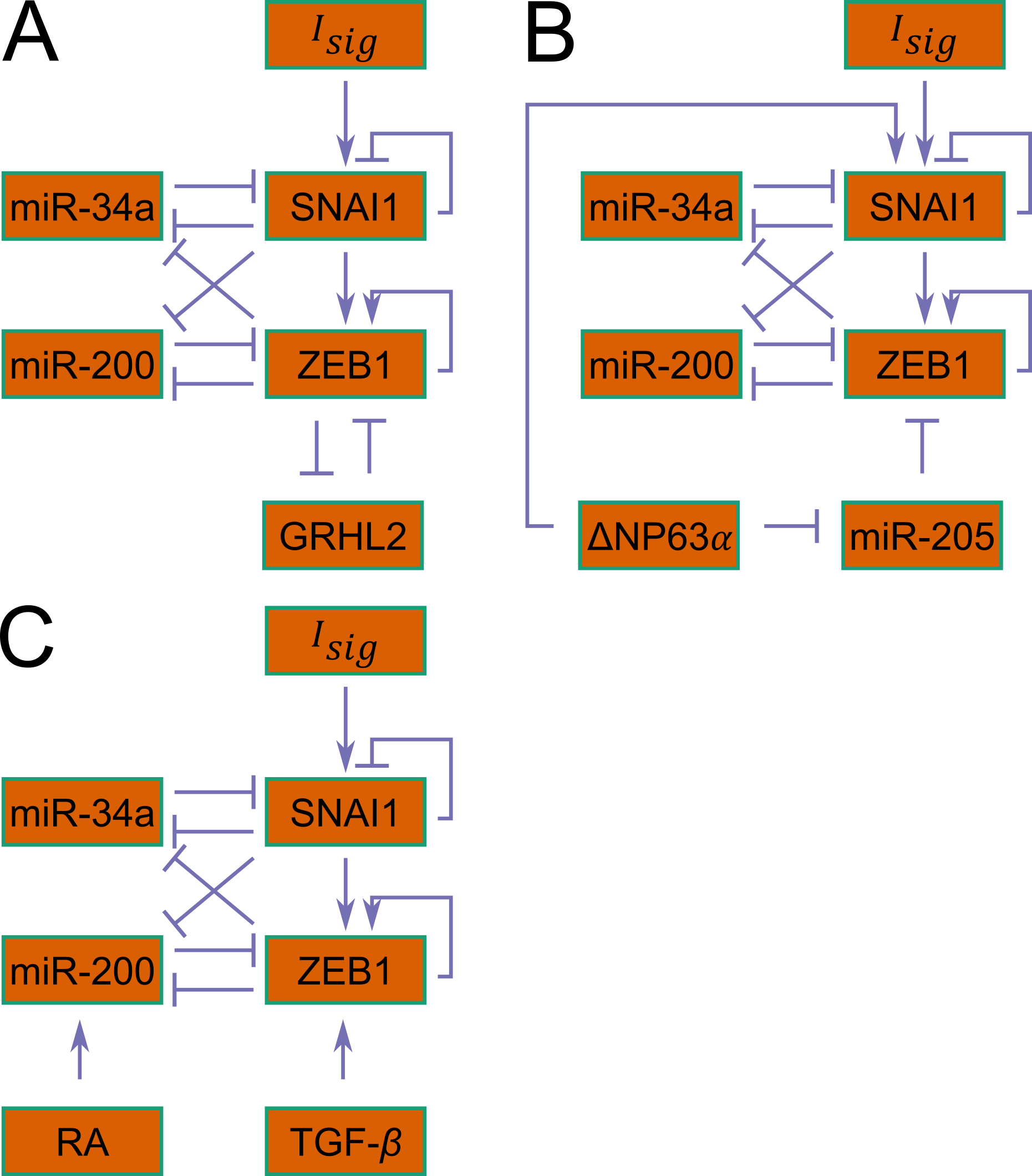

**Fig. SI 1 Coupling of EMT modulators with the regulatory circuit driving EMT and MET. (A)** ZEB1 inhibits the expression of *GRHL2*. GRHL2 inhibits the expression of *ZEB1* (Jolly et al., 2016). **(B)** miR-205 inhibits the expression of *ZEB1*. Expression of miR-205 is inhibited by $\Delta$NP63$\alpha$ which also activates the expression of *SNAI1* (Jolly et al., 2017). **(C)** Retinoic acid (RA) inhibits the activity of miR-200 (Wu et al., 2017) while TGF-$\beta$ promotes the expression of ZEB1 (Gregory et al., 2011; Joseph et al., 2014).

We started the simulation with $500$ cells in the population. The concentration of $I_{sig}$ in each of these cells was drawn from a log-normal distribution with median $2\times{10}^{4}$ molecules / cell and coefficient of variation $1.0$. This choice of the initial distribution of $I_{sig}$ concentrations led to a population which was $\sim84\%$ epithelial, ~$9\%$ hybrid E / M, and ~ $7\%$ mesenchymal. Such a population is representative of the epithelial-mesenchymal heterogeneity characteristic of solid tumors and allowed us to probe the dynamics of phenotypes in cancer cells.

For each cell in the population, we maintained a record of the time point at which the state of the regulatory circuit within the cell had been last updated. When a cell was chosen for division, the concentrations of proteins and RNAs within the cell were updated using the ODEs listed before. Thereafter, the contents of the cell were partitioned among the daughter cells as described in the main text. When a cell was chosen to die, it was deleted from the population. After each cell division or cell death event, the state of the regulatory circuit within each cell in the population was updated using the ODEs listed before. This update was carried out for the period of time between two update events. For example, say the $n^{th}$ cell division / death event occurred at time $t_{n}$. Then, the dynamics of the regulatory circuit within cell $i$ was simulated for the time period $t_{n}-t^{i}$ after carrying out the cell division / cell death event. Here, $t^{i}$ is the last time point at which the state of the regulatory circuit within cell $i$ had been updated.

At each point in time, each cell in the population could be assigned a phenotype based on where the concentrations of *ZEB1* mRNA and $I_{sig}$ within the cell lay with respect to the bifurcation diagram of the EMT / MET regulatory circuit. As shown in fig. 1 (B) in the main text, steady states of the EMT / MET regulatory circuit can be mapped to an epithelial state, a hybrid E / M state, or a mesenchymal state. Each cell in the population was assigned the phenotype of the steady state within the basin of attraction of which it lay.

Drug activity was modeled by increasing the death rate of cells by a multiplicative factor as a function of the drug concentration. In the presence of a drug targeting a specific phenotype, the death rate of cells of that phenotype was given as

$$d=d_{0}(\lambda_{drug}+\frac{1.0-\lambda_{drug}}{1.0+\left( \frac{\left[ D \right]}{\left[ D \right]_{0}} \right)^{n_{drug}}})$$

where $d_{0}$ is the death rate in the absence of the drug, $[D]$ is the drug concentration, $\lambda_{drug}=20.0$ is the maximum possible fold change in the death rate of cells, $n_{drug}=4.0$ is the Hill coefficient describing the pharmacodynamic activity, and $\left[ D \right]_{0}=5.0\times{10}^{4}$ molecules / cell is the threshold drug concentration characterizing drug activity.

Below, we have described, in detail, the simulation regimes used to obtain the figures in the main text.

**Figure 2 (A) and (B)**

Starting with a population of $500$ cells as mentioned above, the simulation was carried out for a period of $8$ weeks. The total carrying capacity ($K$) was fixed at $10000$. While tracking the phenotypes of the parent and the daughter cells during each cell division event (fig. 2 (A)), we used the phenotype of the parent cell right before it divided and the phenotype of the daughter cell right before it divided or died. While plotting the phenotypic composition of the population at a given time point (fig. 2 (B)), we used the phenotype of each cell as determined by the protein and mRNA concentrations at that point in time.

**Figure 2 (C) and (D)**

Starting with a population of $500$ cells as mentioned above, the simulation was carried out for a period of $28$ days. The total carrying capacity ($K$) was fixed at $10000$. This represented *in vivo* growth of tumor cells. Thereafter, we carried out *in silico* FACS to obtain a population of only epithelial cells, only mesenchymal cells, or only hybrid E / M cells. After FACS, we had a phenotypically “pure” population with $100$ cells. In total, we had $3$ such populations- one with only epithelial cells, one with only mesenchymal cells, and one with only hybrid E / M cells. The dynamics of each such population was then simulated for a period of $16$ days with a total carrying capacity ($K)$ of $500$. The lower carrying capacity accounted for the reduced availability of nutrients in the *in vitro* growth conditions.

**Figure 3**

Starting with a population of $500$ cells as mentioned above, the simulation was carried out for a period of $28$ days. The total carrying capacity ($K$) was fixed at $10000$. This represented *in vivo* growth of tumor cells. Thereafter, we carried out *in silico* FACS to obtain a population of $100$ epithelial cells. The dynamics of this population was then simulated for a period of $28$ days with a carrying capacity ($K$) of $500$. From day $1$ to day $10$, $10000$ molecules / cell of $I_{sig}$ were added to the population each day. From day $11$ to day $20$, $10000$ molecules / cell of $I_{sig}$ were subtracted from the population each day. If the $I_{sig}$ concentration in a cell became negative during the withdrawal of $I_{sig}$ dosages, $I_{sig}$ concentration in that cell was set to $0.0$. Day $21$ onwards, there was no external intervention and the dynamics were simulated as described before. Here, we have used a simple simulation setup to understand the population behavior under different external dosage regimes. Our setup does not incorporate the active and passive transport, across cell membranes, of $I_{sig}$ added to the cell culture. These processes are likely to change the time scale over which the effects of different dosages of $I_{sig}$ are observed.

**Figure 4**

To plot fig. 4 (B), the same simulation setup as for fig. 2 (A) was used. To plot fig. 4 (C), the same simulation setup as for fig. 2 (C) was used.

**Figure 5**

Starting with a population of $500$ cells as mentioned above, the simulation was carried out for a period of $28$ days. The total carrying capacity ($K$) was fixed at $10000$. This represented *in* vivo growth of tumor cells. Thereafter, we carried out *in silico* FACS to obtain a population of $100$ epithelial cells. The dynamics of this population was then simulated for a period of $16$ days with a carrying capacity ($K$) of $500$ in the presence of different concentrations of retinoic acid (RA) and TGF-$\beta$.

**Figure 6 and figure 7**

Starting with a population of $500$ cells as mentioned above, the simulation was carried out for a period of $8$ weeks in the absence of drug activity. The dynamics was then simulated for $28$ days in the presence of drug activity. The total carrying capacity ($K$) was fixed at $10000$ for the entire simulation run

For all simulations, we carried out $16$ distinct runs with different seeds for the random number generator. Unless specified otherwise, $\eta=1.9\times{10}^{3}$ was used to obtain the results reported in this study. This value of the noise parameter $\eta$ gave the best fit to the experimental data from Ruscetti *et al*. (Ruscetti et al., 2016) (fig. 2 (C), fig. 2 (D), and fig. S2).

### **Table 1**

| Parameter | Value | Parameter | Value |
| --- | --- | --- | --- |
| $\boldsymbol{g}_{\boldsymbol{\mu}_{\boldsymbol{34}}}$ | $1.35\times{10}^{3} mol.h^{-1}$ | $n_{Z}^{\mu_{34}}$ | $1$ |
| $\boldsymbol{g}_{\boldsymbol{m}_{\boldsymbol{S}}}$ | $90.0 mol.h^{-1}$ | $n_{S}^{m_{S}}$ | $1$ |
| $\boldsymbol{g}_{\boldsymbol{S}}$ | $0.1\times{10}^{3} h^{-1}$ | $n_{I}^{m_{S}}$ | $1$ |
| $\boldsymbol{g}_{\boldsymbol{\mu}_{\boldsymbol{200}}}$ | $2.1\times{10}^{3} mol.h^{-1}$ | $n_{Z}^{\mu_{200}}$ | $3$ |
| $\boldsymbol{g}_{\boldsymbol{m}_{\boldsymbol{Z}}}$ | $11.0 mol.h^{-1}$ | $n_{S}^{\mu_{200}}$ | $2$ |
| $\boldsymbol{g}_{\boldsymbol{Z}}$ | $0.1\times{10}^{3} h^{-1}$ | $n_{Z}^{m_{Z}}$ | $2$ |
| $\boldsymbol{g}_{\boldsymbol{m}_{\boldsymbol{G}}}$ | $22.0 mol.h^{-1}$ | $n_{S}^{m_{Z}}$ | $2$ |
| $\boldsymbol{g}_{\boldsymbol{G}}$ | $200.0 h^{-1}$ | $n_{G}^{m_{Z}}$ | $1$ |
| $\boldsymbol{g}_{\boldsymbol{\mu}_{\boldsymbol{205}}}$ | $2.0\times{10}^{3} mol.h^{-1}$ | $n_{Z}^{m_{G}}$ | $3$ |
| $\boldsymbol{k}_{\boldsymbol{\mu}_{\boldsymbol{34}}}$ | $0.05 h^{-1}$ | $n_{P_{63}}^{m_{S}}$ | $1$ |
| $\boldsymbol{k}_{\boldsymbol{m}_{\boldsymbol{S}}}$ | $0.5 h^{-1}$ | $n_{P_{63}}^{\mu_{205}}$ | $2$ |
| $\boldsymbol{k}_{\boldsymbol{S}}$ | $0.125 h^{-1}$ | $n_{\mu_{205}}^{m_{Z}}$ | $2$ |
| $\boldsymbol{k}_{\boldsymbol{\mu}_{\boldsymbol{200}}}$ | $0.05 h^{-1}$ | $\lambda_{S}^{\mu_{34}}$ | $0.1$ |
| $\boldsymbol{k}_{\boldsymbol{m}_{\boldsymbol{Z}}}$ | $0.5 h^{-1}$ | $\lambda_{S}^{m_{S}}$ | $0.1$ |
| $\boldsymbol{k}_{\boldsymbol{Z}}$ | $0.1 h^{-1}$ | $\lambda_{Z}^{\mu_{34}}$ | $0.2$ |
| $\boldsymbol{k}_{\boldsymbol{m}_{\boldsymbol{G}}}$ | $0.5 h^{-1}$ | $\lambda_{I}^{m_{S}}$ | $10.0$ |
| $\boldsymbol{k}_{\boldsymbol{G}}$ | $0.1 h^{-1}$ | $\lambda_{Z}^{\mu_{200}}$ | $0.1$ |
| $\boldsymbol{k}_{\boldsymbol{\mu}_{\boldsymbol{205}}}$ | $0.05 h^{-1}$ | $\lambda_{S}^{\mu_{200}}$ | $0.1$ |
| $\boldsymbol{S}_{\boldsymbol{0}}^{\boldsymbol{\mu}_{\boldsymbol{34}}}$ | $300.0\times{10}^{3} mol.$ | $\lambda_{Z}^{m_{Z}}$ | $7.5$ |
| $\boldsymbol{S}_{\boldsymbol{0}}^{\boldsymbol{m}_{\boldsymbol{S}}}$ | $200.0\times{10}^{3} mol.$ | $\lambda_{S}^{m_{Z}}$ | $10.0$ |
| $\boldsymbol{Z}_{\boldsymbol{0}}^{\boldsymbol{\mu}_{\boldsymbol{34}}}$ | $600.0\times{10}^{3} mol.$ | $\lambda_{G}^{m_{Z}}$ | $0.65$ |
| $\boldsymbol{\mu}_{\boldsymbol{34}}^{\boldsymbol{0}}$ | $10.0\times{10}^{3} mol.$ | $\lambda_{Z}^{m_{G}}$ | $0.5$ |
| $\boldsymbol{I}_{\boldsymbol{0}}^{\boldsymbol{m}_{\boldsymbol{S}}}$ | $50.0\times{10}^{3} mol.$ | $\lambda_{P_{63}}^{m_{S}}$ | $2.0$ |
| $\boldsymbol{Z}_{\boldsymbol{0}}^{\boldsymbol{\mu}_{\boldsymbol{200}}}$ | $220.0\times{10}^{3} mol.$ | $\lambda_{P_{63}}^{\mu_{205}}$ | $4.0$ |
| $\boldsymbol{S}_{\boldsymbol{0}}^{\boldsymbol{\mu}_{\boldsymbol{200}}}$ | $180.0\times{10}^{3} mol.$ | $\lambda_{\mu_{205}}^{m_{Z}}$ | $0.5$ |
| $\boldsymbol{Z}_{\boldsymbol{0}}^{\boldsymbol{m}_{\boldsymbol{Z}}}$ | $25.0\times{10}^{3}mol.$ | $R_{0}^{\mu_{200}}$ | $1.5\times{10}^{3} mol.$ |
| $\boldsymbol{S}_{\boldsymbol{0}}^{\boldsymbol{m}_{\boldsymbol{Z}}}$ | $180.0\times{10}^{3} mol.$ | $T_{0}^{m_{Z}}$ | $60.0$ |
| $\boldsymbol{\mu}_{\boldsymbol{200}}^{\boldsymbol{0}}$ | $10.0\times{10}^{3} mol.$ | $n_{R}^{\mu_{200}}$ | $1$ |
| $\boldsymbol{G}_{\boldsymbol{0}}^{\boldsymbol{m}_{\boldsymbol{Z}}}$ | $25.0\times{10}^{3} mol.$ | $n_{T}^{m_{Z}}$ | $6$ |
| $\boldsymbol{Z}_{\boldsymbol{0}}^{\boldsymbol{m}_{\boldsymbol{G}}}$ | $10.0\times{10}^{3} mol.$ | $\lambda_{R}^{\mu_{200}}$ | $4.0$ |
| $\boldsymbol{P}_{\boldsymbol{63}_{\boldsymbol{0}}}^{\boldsymbol{m}_{\boldsymbol{S}}}$ | $8.0\times{10}^{3} mol.$ | $\lambda_{T}^{m_{Z}}$ | $35.0$ |
| $\boldsymbol{P}_{\boldsymbol{63}_{\boldsymbol{0}}}^{\boldsymbol{\mu}_{\boldsymbol{205}}}$ | $5.0\times{10}^{3} mol.$ |  |  |
| $\boldsymbol{\mu}_{\boldsymbol{205}_{\boldsymbol{0}}}^{\boldsymbol{m}_{\boldsymbol{Z}}}$ | $10.0\times{10}^{3} mol.$ |  |  |
| $\boldsymbol{n}_{\boldsymbol{S}}^{\boldsymbol{\mu}_{\boldsymbol{34}}}$ | $1$ |  |  |

### **Table 2**

| No. of miRNA binding sites | 0 | 1 | 2 | 3 | 4 | 5 | 6 |
| --- | --- | --- | --- | --- | --- | --- | --- |
| $\boldsymbol{l}_{\boldsymbol{i}}$ | $1.0$ | $0.6$ | $0.3$ | $0.1$ | $0.05$ | $0.05$ | $0.05$ |
| $\boldsymbol{\gamma}_{\boldsymbol{m}_{\boldsymbol{i}}}\boldsymbol{(}\boldsymbol{h}^{\boldsymbol{-1}}\boldsymbol{)}$ | $0.0$ | $0.04$ | $0.2$ | $1.0$ | $1.0$ | $1.0$ | $1.0$ |
| $\boldsymbol{\gamma}_{\boldsymbol{\mu}_{\boldsymbol{i}}}\boldsymbol{(}\boldsymbol{h}^{\boldsymbol{-1}}\boldsymbol{)}$ | $0.0$ | $0.005$ | $0.05$ | $0.5$ | $0.5$ | $0.5$ | $0.5$ |

Here, $mol.\equiv molecules / cell$.
