## Supplementary Figures for "A Mechanism for Epithelial-Mesenchymal Heterogeneity in a Population of Cancer Cells"

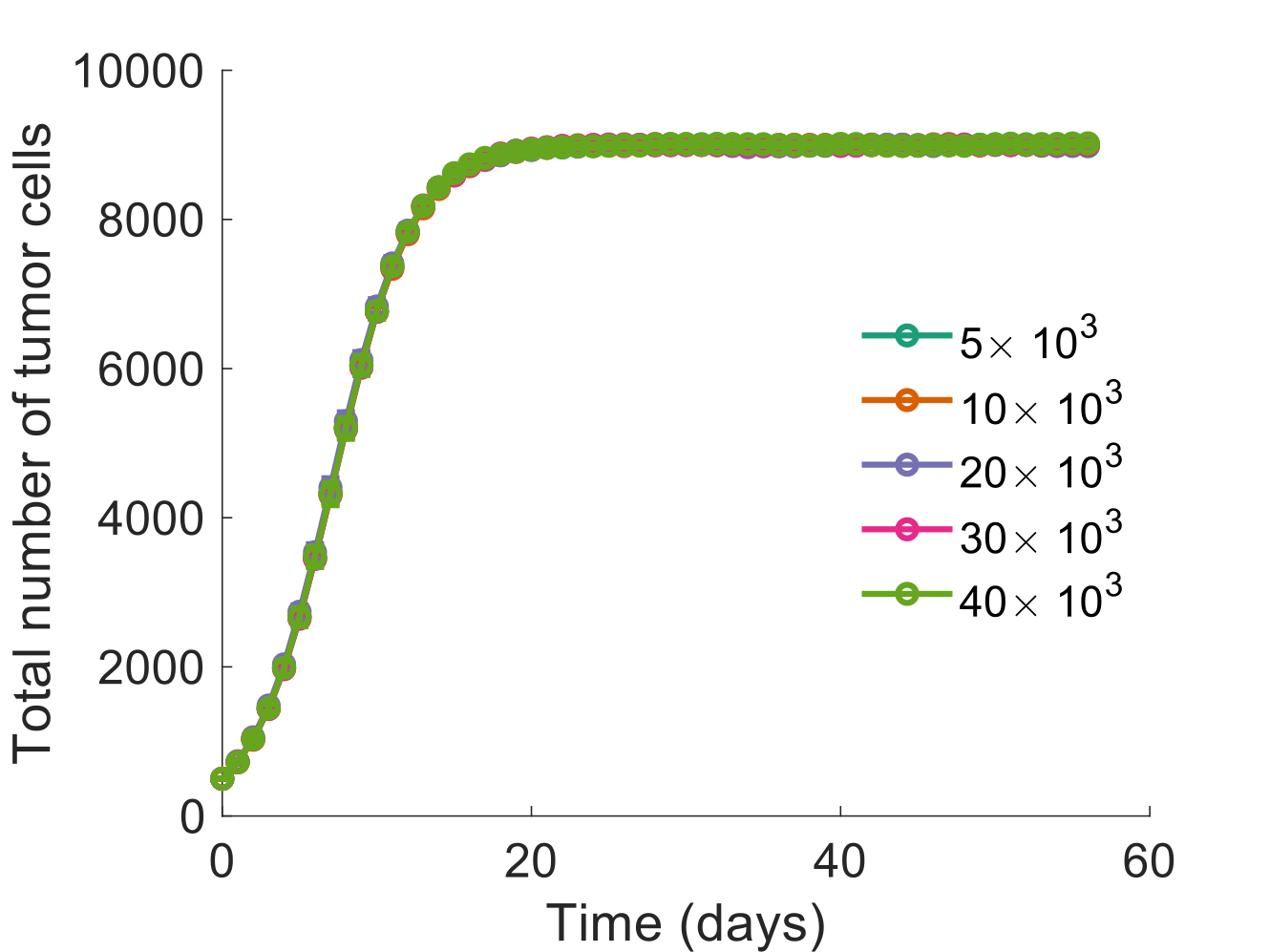

**Figure S1** **Growth kinetics of tumor cells for different values of the noise parameter** $\boldsymbol{\eta}$**.** Simulations were started with a population of $500$ cells on day $0$ and a fixed carrying capacity of $10000$ cells. In each case, the number of tumor cells became nearly stationary around day $18$, independent of $\eta$. The number of tumor cells at different points shown here was obtained by averaging over $16$ distinct simulation runs. Error bars indicate the standard deviation in the number of tumor cells at each time point calculated over these runs.

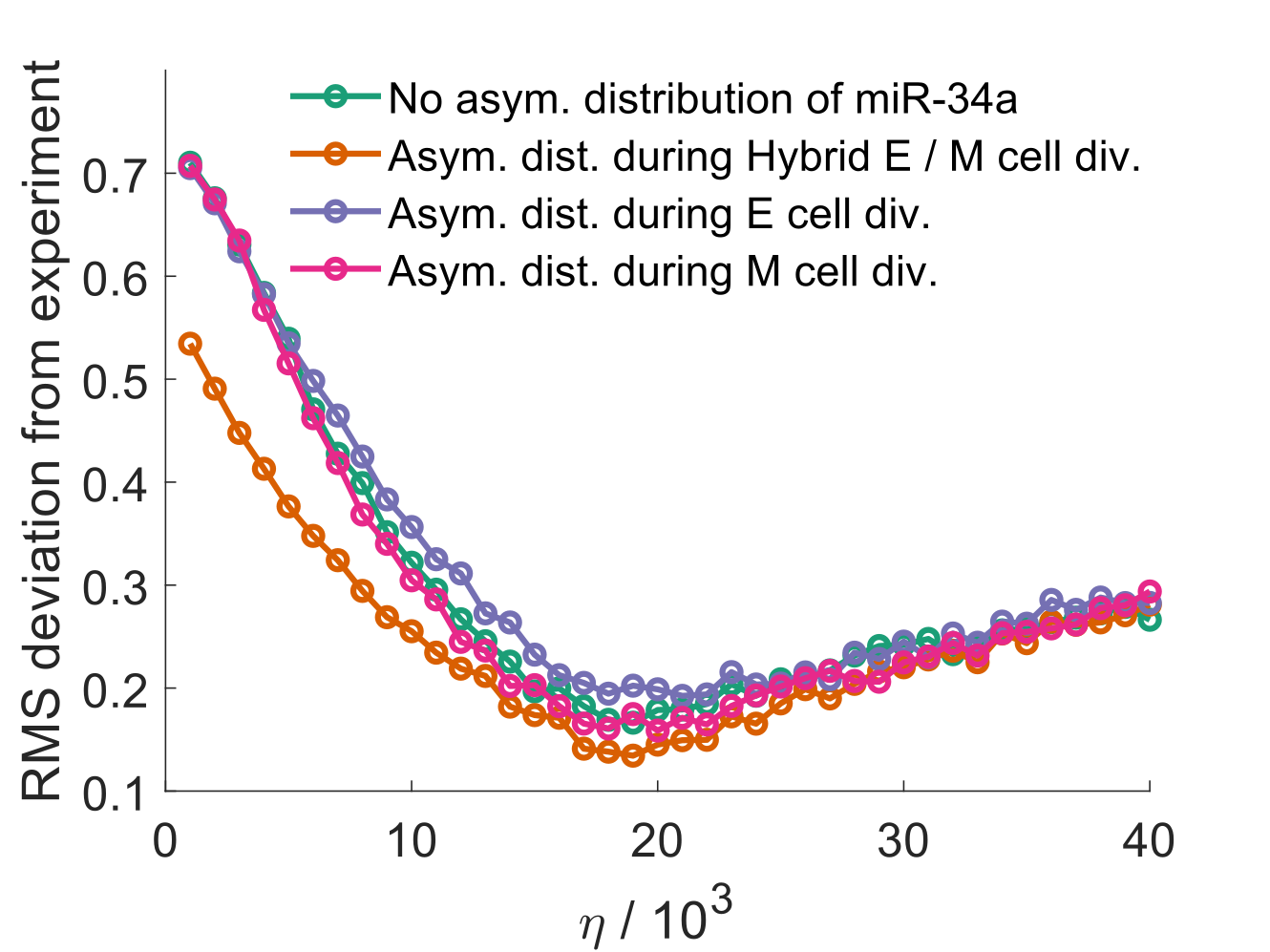

**Figure S2** **Root mean square deviation (RMSD) of model predictions from experimental data for murine prostate cancer cells obtained from Ruscetti *et al.*** RMSD was calculated by pooling fractions of the three phenotypes at different time points obtained in three cases— when starting with a population of only epithelial cells, when starting with a population of only hybrid E / M cells, and when starting with a population of only mesenchymal cells. A lower RMSD was obtained in the model with asymmetric distribution of miR-34a among the daughter cells during the division of hybrid E / M cells.

$I_{sig}$ concentration above a threshold leads to a mesenchymal phenotype. Also, this concentration cannot fall before $0.0$. Thus, at very high values of $\eta$, the fraction of mesenchymal cells in the population will increase rapidly leading to large deviation from experimental data at high $\eta$. At low $\eta$, the probability of a daughter cell acquiring a phenotype different from that of the parent cell will be very low due to only a small change in $I_{sig}$ concentration during cell division. Thus, at very low values of $\eta$, the model will not be able to capture the plasticity of hybrid E / M populations. Large deviations from experimental behavior at low and high $\eta$ account for the non-monotonic nature of the curves in this figure.

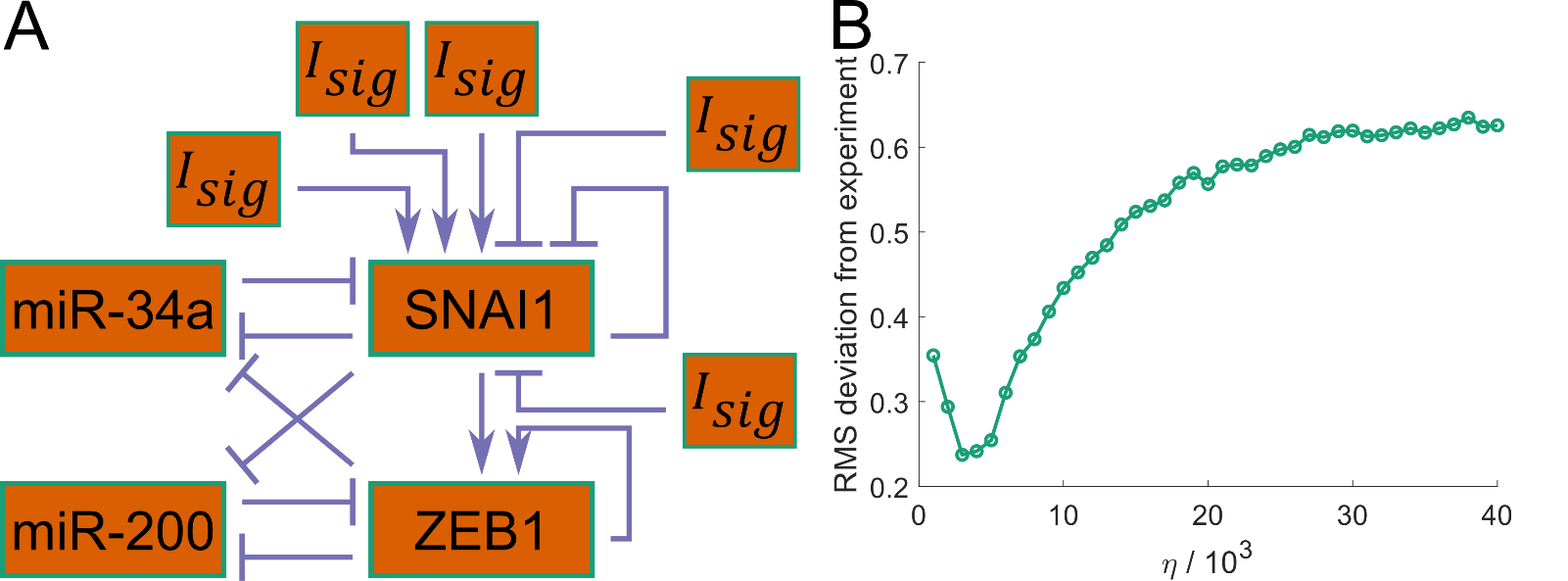

**Figure S3** **Fitting experimental data in the presence of multiple signaling pathways that converge onto the EMT / MET regulatory circuit.** These include signals promoting EMT and those inhibiting EMT. We simulated the proposed EMP model while considering these signals independently. **(A)** The EMT / MET regulatory circuit with $3$ EMT-inducing and $2$ EMT-inhibiting signals. **(B)** Root mean square deviation (RMSD) of model predictions from experimental data for murine prostate cancer cells obtained from Ruscetti *et al.* The model predictions were obtained using the regulatory circuit shown in (A) with the same value of the noise parameter $\eta$ for each input signal. Panel (B) indicates that, in the presence of multiple EMT-inducing and EMT-inhibiting signals, a good fit to experimental data can be obtained at a much lower value of the noise parameter $\eta$. The kinetic parameters governing the regulation of *SNAI1* by $I_{sig}$ was kept the same for each input to the regulatory circuit. Only $\lambda_{I}^{m_{S}}$, the fold change in the production rate of *SNAI1* mRNA in response to the input, differed for the activating and inhibitory inputs. For activating inputs, $\lambda_{I}^{m_{S}}$=$10.0$, and for inhibitory inputs, $\lambda_{I}^{m_{S}}$=$0.1$.

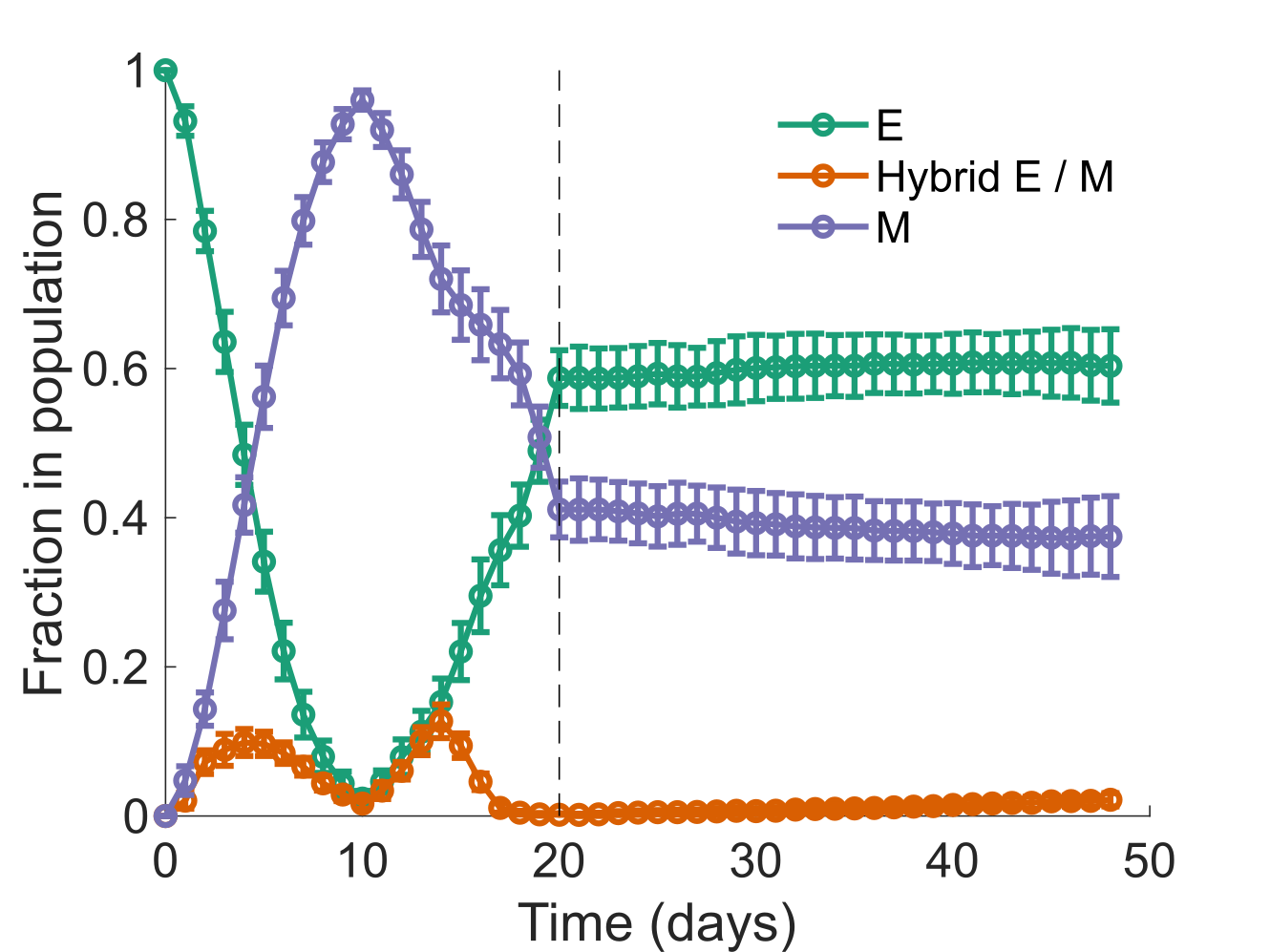

**Figure S4** **Retention of the mesenchymal phenotype after the EMT-inducing signal has been withdrawn.** Starting with a population of epithelial cells on day $0$, ${10}^{3}$ molecules / cell of $I_{sig}$ was added to the population each day till day $10$. Starting on day $11$, the same concentration of $I_{sig}$ was removed from the population each day. After day $20$ (indicated by the black dashed line), no addition or withdrawal of $I_{sig}$ dosages was administered. The fraction of epithelial and mesenchymal cells in the population on day $20$ was maintained over the next many days. $\sim40\%$ of cells in the population underwent an EMT which was irreversible on the time scale considered here. Error bars indicate the standard deviation calculated over $16$ independent runs.

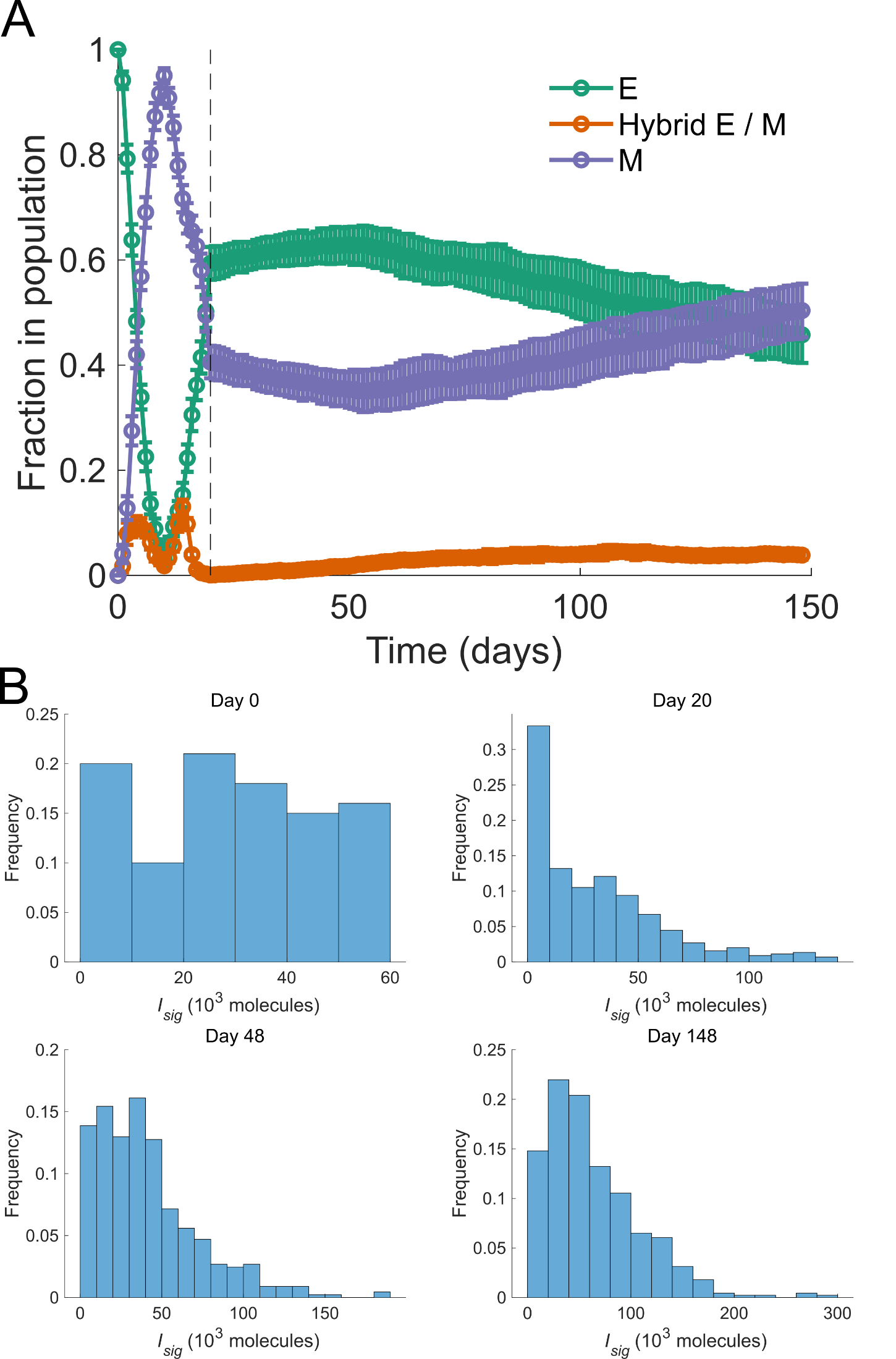

**Figure S5 Long term behavior of the population after the EMT-inducing signal has been withdrawn.** **(A)** Fraction of epithelial, mesenchymal, and hybrid E / M phenotypes over a period of $148$ days in the population considered in fig. S4. The black dashed line indicates day $20$ after which no addition or subtraction of $I_{sig}$ dosages was administered. Error bars indicate the standard deviation calculated over $16$ independent runs. **(B)** Distribution of $I_{sig}$ concentration in cells in the population at different time points during the simulation run.

As shown in (B), at the end of $20$-day period, most cells in the population either have $I_{sig}$ concentration close to $0.0$ or very high $I_{sig}$ concentration. Few cells have $I_{sig}$ concentration in the region of tri-stable dynamics of the EMT regulatory circuit (fig. 1 (B)). Since cells far away from the tri-stable region, which form the bulk of the population on day $20$, are highly unlikely to generate a daughter cell with a phenotype different from that of the parent cell, the fractions of cells of different phenotypes in the population does not change much for nearly a month after day $20$ when the external addition or withdrawal of $I_{sig}$ dosages was stopped. The population takes a long time to recover from the clustering of $I_{sig}$ concentrations away form the tri-stable region. However, noise in the partitioning of $I_{sig}$ among the daughter cells during cell division eventually drives the $I_{sig}$ concentration in cells back into the tri-stable region and around day $57$, we once again start observing the typical dynamics— decrease in the fraction of epithelial cells and increase in the fraction of mesenchymal cells in the population.
